# Endothelial CXCL5 negatively regulates myelination and repair after white matter stroke

**DOI:** 10.1101/664953

**Authors:** Guanxi Xiao, Rosie Kumar, Yutaro Komuro, Jasmine Burguet, Visesha Kakarla, Ida Azizkhanian, Sunil A. Sheth, Christopher K. Williams, Xinhai R. Zhang, Michal Macknicki, Andrew Brumm, Riki Kawaguchi, Phu Mai, Naoki Kaneko, Harry V. Vinters, S. Thomas Carmichael, Leif A. Havton, Charles DeCarli, Jason D. Hinman

## Abstract

Cerebral small vessel disease and resulting white matter pathologies are worsened by cardiovascular risk factors including obesity. The molecular changes in cerebral endothelial cells caused by chronic cerebrovascular risk factors remain unknown. We developed a novel approach for molecular profiling of chronically injured cerebral endothelial cells using cell-specific translating ribosome affinity purification (RiboTag) with RNA-seq in Tie2-Cre:RiboTag mice. We used this approach to identify the transcriptome of white matter endothelial cells after the onset of diet-induced obesity (DIO). DIO induces an IL-17B signaling pathway that acts on the cerebral endothelia through IL-17Rb to increase levels of both circulating CXCL5 and local endothelial expression of CXCL5 in both the DIO mouse model and in humans with imaging or pathologic evidence of cerebral small vessel disease. In the white matter, endothelial CXCL5 acts as a chemoattractant and promotes the association of oligodendrocyte progenitor cells (OPCs) with cerebral endothelia increasing vessel-associated OPC cell number and triggers OPC gene expression programs regulating migration and chemokine receptor activation. Targeted blockade of IL-17B with peripheral antibody administration reduced the population of vessel-associated OPCs by reducing endothelial CXCL5 expression. CXCL5-mediated sequestration of OPCs to white matter vasculature impairs OPC differentiation after a focal white matter ischemic lesion. DIO promotes a unique white matter endothelial-to-oligodendrocyte progenitor cell signaling pathway that compromises brain repair after stroke.

## Introduction

Cerebral small vessel disease is an age-related entity affecting brain white matter. The resulting white matter lesions accumulate over time (1) and contribute to disability (2), dementia (3–5), and death (6). Cerebral small vessel injury is significantly worsened by chronic cardiovascular risk factors such as hypertension, diabetes, and obesity (7–10). In particular, abdominal obesity and its associated metabolic disturbances in blood pressure, lipids, and blood sugar control increase the risk of developing white matter lesions on MRI (11–14) and increase the likelihood of lacunar brain infarction or stroke (15). While the pathologic changes associated with cerebral small vessel disease are well known (16, 17), the molecular pathways that drive small vessel injury in the brain are largely unknown.

Emerging data suggests that an interaction between cerebral vessels and cells of the oligodendrocyte lineage play a key role in maintaining white matter homeostasis (18–20). A subset of platelet-derived growth factor receptor alpha-positive (PDGFRα+) oligodendrocyte progenitor cells (OPCs) closely associate with the vasculature (21) and use it to migrate in the brain during development (22). Proteins secreted by endothelial cells promote OPC migration and proliferation *in vitro* (23, 24). In the spontaneously hypertensive rat model of cerebral small vessel disease, the OPC population is increased in association with vascular changes and delays in OPC maturation may be mediated by endothelial secretion of *HSP90*α (25). Both the diagnosis and treatment of cerebral small vessel disease would be advanced by identifying additional molecular pathways active in cerebral endothelia and driven by chronic cardiovascular risk factors (26).

To identify the molecular changes in white matter endothelia in the setting of chronic cardiovascular risk factors, we used a mouse model of diet-induced obesity (DIO) (27) that recapitulates a number of features of human cardiovascular risk (28). We demonstrate that DIO is associated with a loss of white matter vasculature, increases in the number of OPCs in brain white matter, thinner myelin, and disrupted axons. We used cell-specific translating ribosome affinity purification and RNA-sequencing in Tie2-Cre:RiboTag mice to isolate the endothelial-specific transcriptome after the onset of DIO. This approach led to the identification of a novel IL-17 signaling cascade acting through the IL-17B/IL-17Rb isoforms of the IL-17 family that is active in chronically injured cerebral endothelial cells and increases endothelial expression of CXCL5. Endothelial over-expression of CXCL5 directly signals to OPCs acting as a chemoattractant *in vivo* and triggers OPC gene expression programs regulating migration and chemokine receptor activation. DIO-induced endothelial expression of this immune signaling pathway exacerbates the white matter injury response to a focal white matter ischemic lesion and restricts the maturation of OPCs during the repair phase after stroke. Increased serum levels of CXCL5 are found in a subset of human subjects at risk for cerebrovascular disease and within cerebral endothelial cells of individuals with cerebrovascular damage. These findings provide evidence that chronic cerebrovascular risk factors can promote vascular regulation of myelination and have direct implications for the understanding of human cerebral small vessel disease and repair of cerebral white matter.

## Results

### Diet-induced obesity as a model of white matter and vascular injury

Obesity is a significant risk factor for the development of small vessel disease and white matter injury (10, 11, 13, 14). We used a well-established model of diet-induced obesity (DIO) (27) to model the effects of chronic cardiovascular risk on brain white matter and the vasculature. Beginning at 8 weeks of age, mice were fed a control fat diet (CFD) or a high fat diet (HFD) for 12 weeks. After 12 weeks on the dietary intervention, mice exhibit 84% weight gain and metabolic disturbances in cholesterol and blood sugar (Fig. S1) broadly consistent with the diagnostic criteria for metabolic syndrome (29). After the development of obesity, we examined the vasculature and cellular makeup of the white matter.

Using a Tie2-Cre;tdTomato (Ai14) strain that robustly labels the vasculature throughout the brain, we characterized the effects of DIO on the white matter vasculature throughout the corpus callosum (Fig. 1A). DIO reduces the volume of tdT+ vessels by 26.0% and the branch complexity of the vasculature (Fig. 1B). DIO also resulted in an increase in the PDGFRα+ oligodendrocyte progenitor cells (OPCs) within the corpus callosum and a concordant increase in OPCs associated with vessels, measured as OPCs per unit vessel length (Fig. 1C-D). Consistent with a DIO-induced increase in OPCs, we observed fewer and shorter axonal paranodal segments (Fig. S2A) as well as thinner myelin with an increased *g*-ratio as measured by electron microscopy (Fig. S2B), indicating compromise of white matter integrity in DIO mice. We developed a direct RNA hybridization assay for oligodendrocyte staging based on overall marker gene expression patterns in white matter, clustered by the three main stages of oligodendrocyte development. The top 40 genes marking OPCs, pre-myelinating oligodendrocytes (PMO), and myelinating oligodendrocytes (MO) (30) were used to indicate oligodendrocyte stages in mice on Using this 120 gene expression platform (SI Data File 1; Fig. 1E, Fig. S2C), we found an increase in OPC gene expression in DIO white matter, with animals on HFD clustering more closely with genetically defined OPCs, consistent with light microscopy findings. Together, these findings suggested that DIO increases OPC proliferation, increases the number of OPCs associated with vessels, and biases the oligodendrocyte lineage towards immaturity.

**Figure 1.**
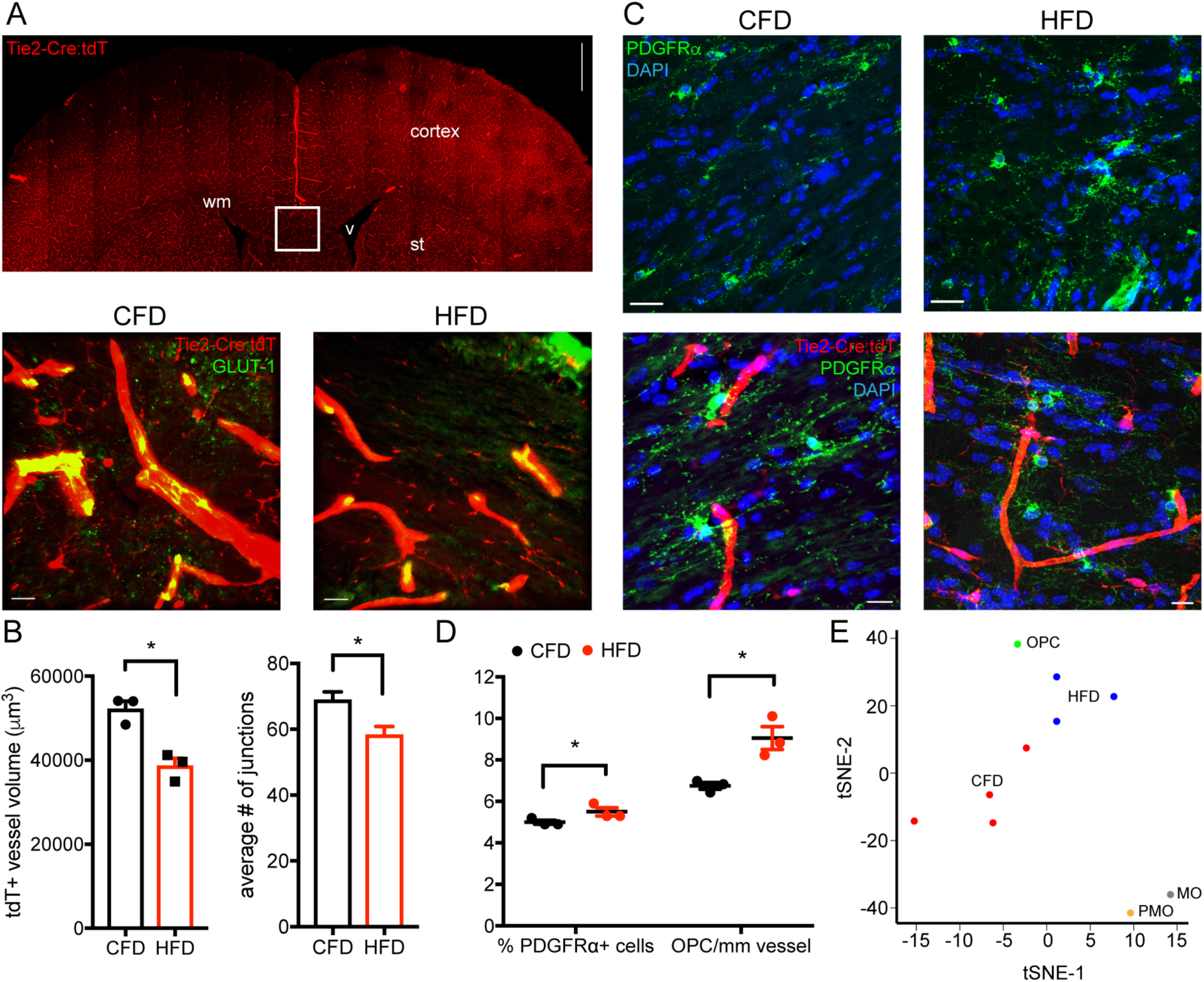
Diet-induced obesity as a model of white matter and vascular injury. Tie2-Cre;tdTomato transgenic mice (*n*=3/grp; 15 confocal z-stacks per animal; inset box) were used to measure vascular changes after DIO (A). Images of white matter from CFD (left lower panel) and HFD (right lower panel) animals labeled for Tie2-Cre;tdTomato (red) and GLUT-1 (green). Average white matter tdT+ vessel volume in CFD (black) and HFD (red) animals (*p*=0.0069; left) and average vascular junctions (*p*=0.0032; right) (B). PDGFRα+ OPCs (green) in CFD (left upper panel) and HFD (right upper panel) white matter. Vessel-associated PDGFRα+ OPCs in CFD (left lower panel) and HFD (right lower panel) (C). Percentage of PDGFRα+ OPCs is increased in HFD (5.01±0.13% vs. 5.66±0.22%; \**p*=0.014) and the number of PDGFRα+ OPCs per mm vessel length in increased (6.46±0.23 vs. 8.94±0.31 cells/mm; \**p*<0.0001) (D). tSNE of Nanostring gene expression for CFD (red, *n*=4) and HFD (blue, *n*=3) animals using reference profiles of OPCs, pre-myelinating oligodendrocytes (PMO), and myelinating oligodendrocytes (MO). Scale bars = 10 μm.

### Molecular profiling of white matter endothelia

Systemic cardiovascular risk factors such as DIO exert their effect on white matter by primarily damaging the cerebrovasculature. To identify the molecular pathways induced in white matter endothelia, we developed an approach using Tie2-Cre:RiboTag mice together with translating ribosome affinity purification using the RiboTag method (31) (Fig. 2A). This transgenic approach leads to robust HA labeling in the cerebrovasculature (Fig. 2B). HA-immunoprecipitated RNA using this approach shows endothelial specificity, with a specific enrichment of endothelial transcripts (Fig. 2C) compared to established marker genes for other perivascular cells including pericytes and OPCs (30). DIO results in a specific gene expression profile compared to endothelial cells from normal weight mice (Fig. S3A; SI Data File 2) within white matter endothelia: 112 genes are up-regulated and 60 genes are down-regulated (FDR<0.1, Fig. 2D). Among the top differentially regulated genes, *interleukin-17 receptor b* (*IL17Rb*) (8.83-fold increased, FDR=0.090) and its effector chemokine *Cxcl5* (11.35-fold increased, FDR=0.064) were two of the most strongly up-regulated genes when comparing DIO vs. control animals (Fig. 2E). To confirm RNA-seq results, we performed TRAP-qPCR using independent Tie2-Cre:RiboTag biologic replicates for a subset of differentially regulated genes (*Glut-1, Itgb3*, *Cd180, Hsd3b3, Tnfrsf10b, Il17rb, Cxcl5, and Ttc21a* ) (Fig. S3B). Similar degrees of up-regulation for *Il17rb* and *Cxcl5* using qPCR were seen. Gene ontology of the up-regulated genes suggests enrichment of immune signaling pathways including C-X-C chemokine signaling and interleukin receptor activation (Fig. S3C).

**Figure 2.**
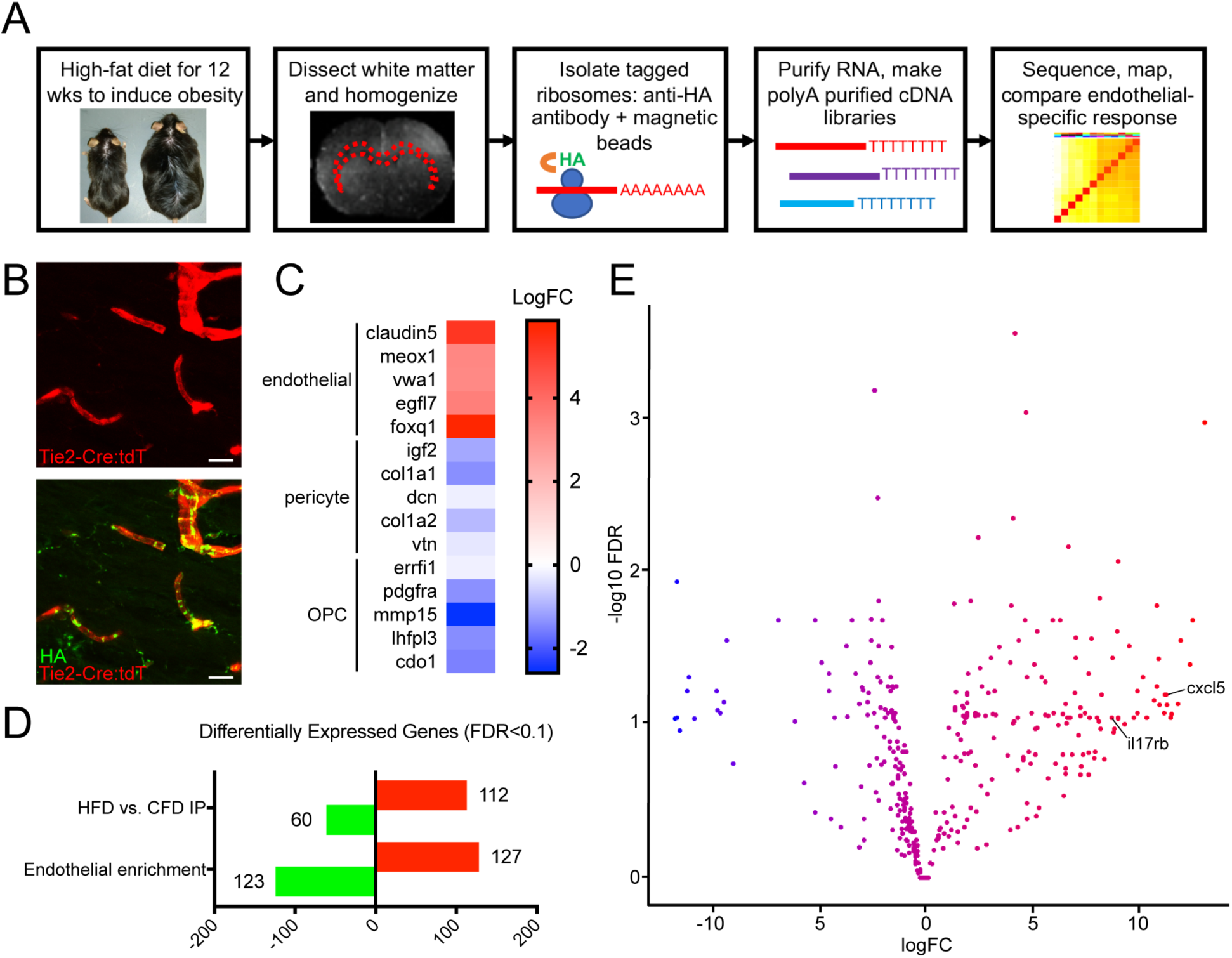
Molecular profiling of white matter endothelia. Schematic representation of workflow for identifying the molecular profile of chronically injured white matter endothelia using translating ribosome affinity purification after DIO (A). Tie2-Cre;tdTomato;RiboTag transgenic mice labeled for HA (B). Anti-HA pulldowns from control white matter show enrichment for endothelial marker genes and de-enrichment of pericyte and OPC markers (C). Differentially expressed genes (FDR<0.1) between anti-HA pulldowns in CFD and HFD animals (*n*=3/grp) and IP vs. input in CFD animals showing the number of genes up- and down-regulated in endothelial in normal weight mice (D). Volcano plot of the top differentially expressed genes between anti-HA pulldowns in CFD and HFD with *Il17rb* and *Cxcl5* labeled. Scale bars = 10 μm. Complete gene list available in SI Data File 2.

### IL-17Rb and CXCL5 up-regulation in human and murine white matter vasculature

IL-17 signaling involves five interleukin ligands (A-E) and five cognate receptor isoforms that hetero and/or homo-dimerize to effect downstream signaling (32). Within our transcriptional dataset, the only IL-17 receptor isoform that was significantly differentially regulated in DIO-affected cerebral endothelial cells was IL-17Rb (SI Data File 2). Among a number of diverse functions, IL-17 receptor activation drives effector chemokine signaling, including CXCL5 (33) as a mechanism of identifying tissue injury. CXCL5 is a member of the C-X-C chemokine family (34) that acts as a chemoattractant in other tissues and has been reportedly up-regulated in white matter after peri-natal hypoxia (35). Guided by our RNA-seq data, we hypothesized that DIO may induce IL-17B signaling acting through IL-17Rb resulting in increased endothelial expression of CXCL5 (Fig. 3A). In primary human brain microvascular endothelial cells, stimulation with recombinant isoforms of IL-17 (A-E) increased the secretion of CXCL5 in conditioned medium with IL-17B, D and E driving two-fold increases in CXCL5 secretion (Fig. 3B). Retro-orbital venous blood sampling confirmed increased serum detection of CXCL5 in DIO mice (Fig. 3C). Immunofluorescent labeling for IL-17Rb (Fig. 3D) and CXCL5 (Fig. 3E) in Tie2-Cre;tdTomato (Ai14) mice demonstrated a marked increase in detection of both molecules within white matter cerebral vessels in DIO mice.

**Figure 3.**
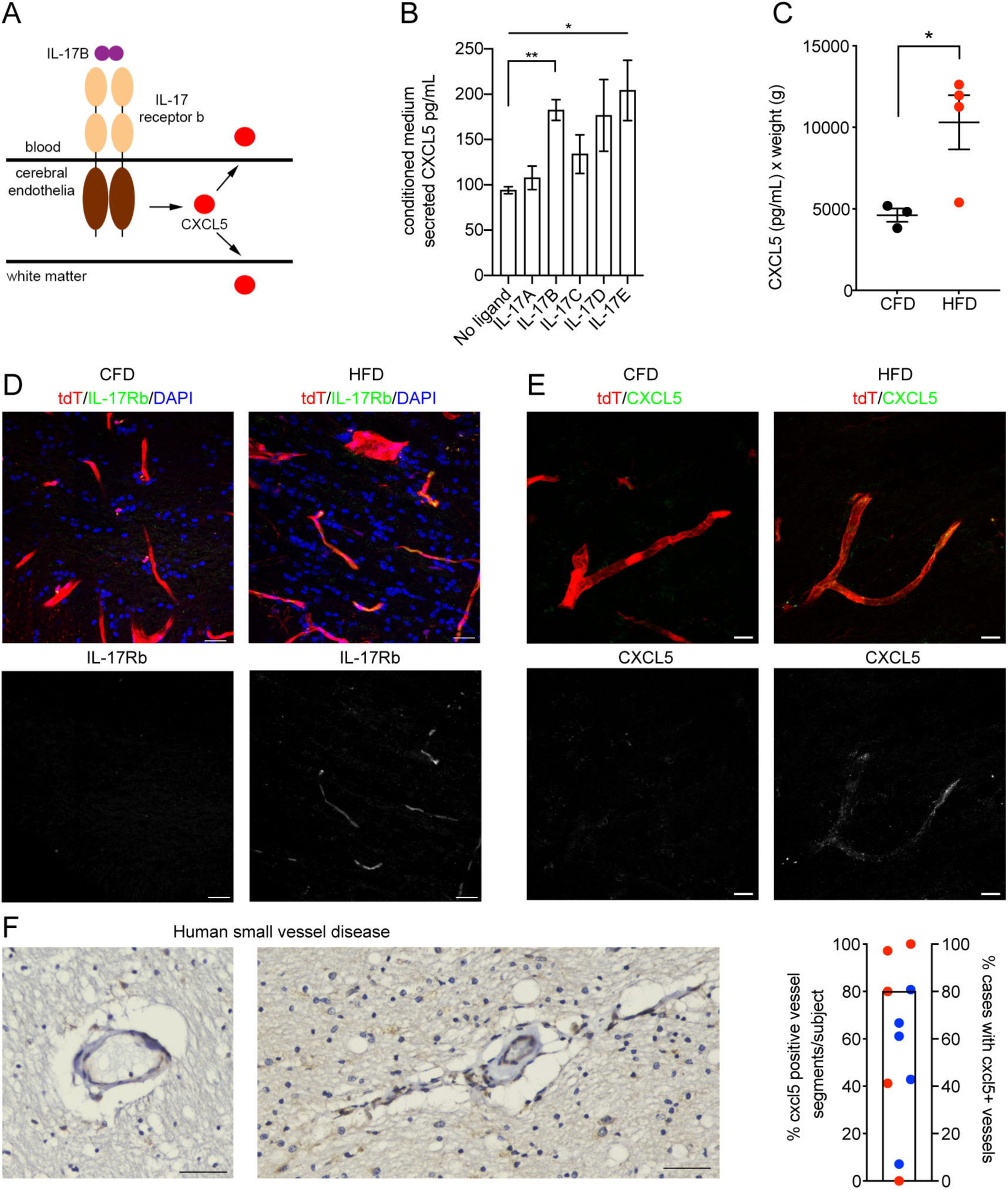
IL-17Rb and CXCL5 up-regulation in injured white matter vasculature. Schematic representation of IL-17/CXCL5 signaling in chronically injured cerebral endothelia (A). Human brain microvascular endothelial cells were stimulated with IL-17 ligands A-E (250 ng/mL) and CXCL5 levels measured in conditioned media 48 hrs after stimulation (\**p*=0.0372 by Kruskal-Wallis H test; **post-hoc comparison for IL-17B vs. no ligand, adjusted *p*=0.0178) (B). Weight adjusted-ELISA values (pg/mL) for murine CXCL5 in retro-orbital blood samples from CFD (black) and HFD (red) animals (*n*=4/grp, *p*=0.0355) (C). Immunofluorescence labeling for IL-17Rb (green, D) and CXCL5 (green, E) is absent in white matter vasculature of Tie2-Cre;tdTomato mice on CFD (left panels) and abundant in white matter vasculature of Tie2-Cre;tdTomato mice on HFD (right panels). Single channel labeling for IL17Rb (lower panels, D) and CXCL5 (lower panels, E) show heterogenous endothelial expression. CXCL5 is detected in human frontal white matter endothelia from aged subjects. Bar represents percentage of cases (8/10) with any CXCL5 staining with points indicating the individual percentage of CXCL5-positive vessel segments in individuals with (blue) and without (red) vascular dementia (F). Scale bars = 50 μm in F, 20 μm in D, 10 μm in E.

To verify the relevance of this DIO-induced cerebrovascular molecular pathway to human cerebral small vessel disease, we examined endothelial CXCL5 expression in a series of older (86±8 years of age) human post-mortem specimens with (*n* =5) and without (*n* =5) a pathologic diagnosis of cerebral vascular disease sufficient to influence cognition in the setting of low levels of Alzheimer’s disease pathology. Sections containing frontal peri-ventricular white matter were immunolabeled for CXCL5 (Fig. S4). In this older cohort, CXCL5 is robustly detected in cerebral vessel segments within white matter, with 80% demonstrating at least some CXCL5 staining within white matter vasculature, while the mean percentage of vessel segments showing CXCL5 staining was 71.2±0.08% (17.2±3.4 vessel segments/subject) (Fig. 3F).

### The IL-17/CXCL5 pathway as a novel vessel-to-OPC signaling paradigm

With the known role of chemokine receptor (CXCR) signaling on OPC migration (22), we reasoned that endothelial up-regulation of CXCL5 in DIO mice may function to promote OPC migration to the vasculature. In callosal white matter, we observed a notable increase in OPCs that were in close apposition to CXCL5+ vessel segments (Fig. 4A) in DIO mice. *In vitro* exposure of O4+ OPCs to increasing doses of recombinant murine CXCL5 resulted in a dose-dependent increase in OPC cell area with cytoskeletal changes suggesting motility (Fig. 4B). To determine the ability of endothelial CXCL5 to signal to OPCs *in vivo*, we used a combined transgenic and targeted viral gene expression approach (Fig. 4C). We designed a pCDH-FLEX-CXCL5-T2A-GFP lentiviral construct and injected this virus or control virus (pCDH-FLEX-GFP) into the subcortical white matter of Tie2-Cre;tdTomato mice resulting in targeted gene expression specifically in white matter vasculature (Fig. S5). After 6 weeks of endothelial up-regulation of CXCL5-GFP or GFP in normal weight mice, we measured the distance of individual OPCs from vessels and the cell area of vessel-associated OPCs, as well as the number of OPCs per unit vessel length (Fig. S6). The average distance of OPCs from tdT+ vessels was reduced in CXCL5-GFP injected animals compared to GFP injected animals while the number of PDGFRα+ OPCs in apposition to tdT+ vessels were increased (upper panels Fig. 4C, 4E, 4F) supporting a chemoattractant role for CXCL5 on OPCs. Similar to the effects of recombinant CXCL5 on OPCs *in vitro*, endothelial over-expression of CXCL5 *in vivo* resulted in increased OPC cell area (lower panels Fig. 4C, 4G).

**Figure 4.**
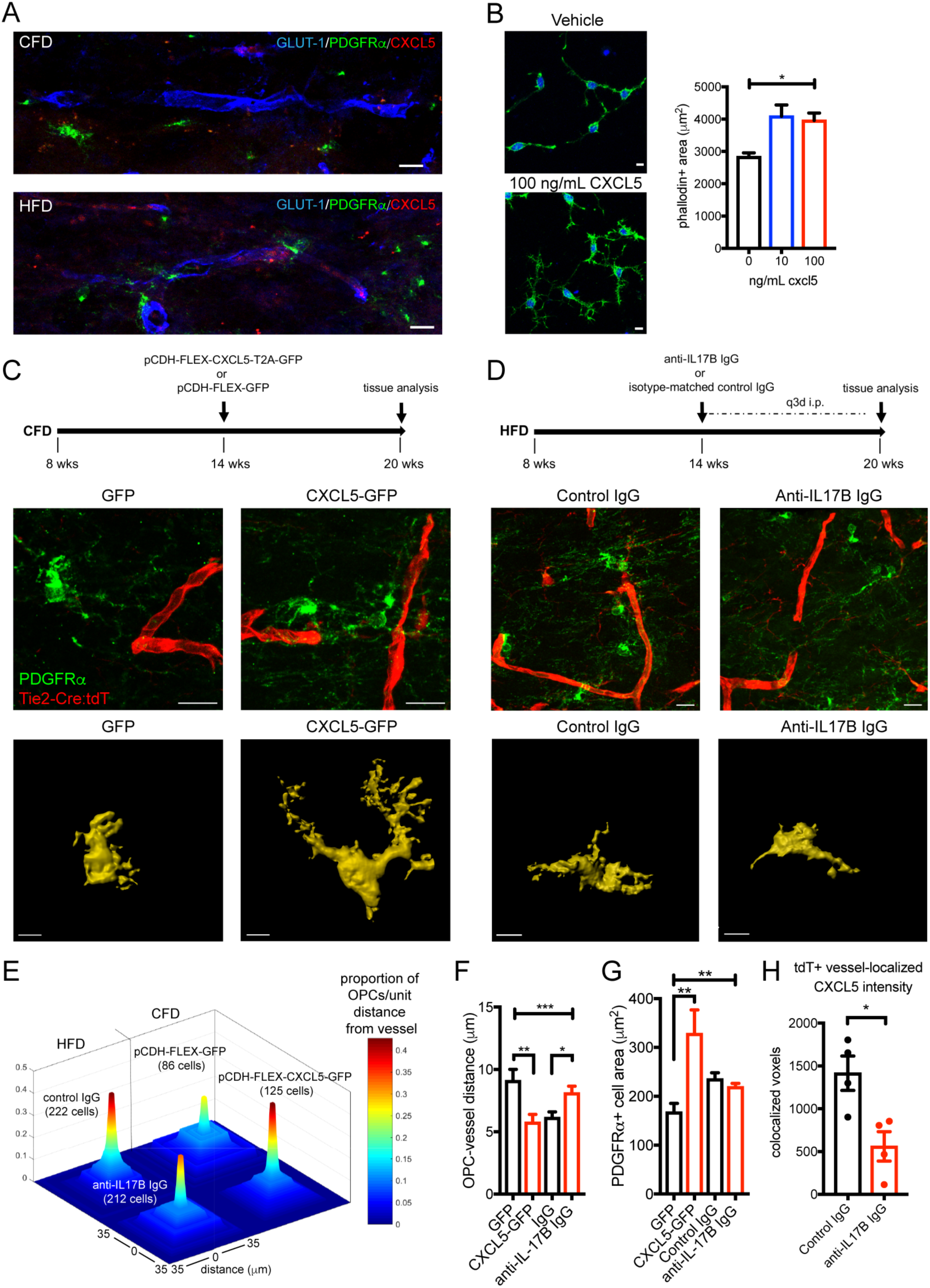
CXCL5 is a novel vessel-to-OPC signal in white matter vasculature. Association of PDGFRα+ OPCs (green) with the vasculature (blue) in CFD (upper panel) and CXCL5-positive (red) HFD animals (lower panel) (A). Phalloidin-positive cellular area in O4+ OPCs grown in vitro exposed to vehicle (upper panel) or recombinant CXCL5 (lower panel) for 48 h (*p*<0.0001, F=9.82 by one-way ANOVA). Experimental approach for CXCL5 transgenic-viral gain of function (upper panel, C). PDGFRα+ OPC (green) labeling in GFP-transduced Tie2-Cre;tdTomato mice (red, left panel) and CXCL5-GFP-transduced Tie2-Cre;tdTomato mice (right panel). Representative masked cellular profiles of PDGFRα+ cell area (lower panels). Schematic of anti-IL-17B antibody treatment (upper panel, D). PDGFRα+ OPC (green) labeling in control IgG-treated Tie2-Cre:tdT mice (left panel) and anti-IL-17B IgG-treated Tie2-Cre:tdT mice (right panel). Representative masked cellular profiles of PDGFRα+ cell area (lower panels). Proportion of OPCs per unit distance from vessel (0-35 μm) in each condition (total measured cell number per condition in parentheses) (E). Average distance of OPCs to vessel (\*\*\**p*=0.0005, F=6.06 by one-way ANOVA; **adjusted *p*=0.0039; *adjusted *p*=0.0168) (F). Average *in vivo* PDGFRα+ OPC cell area (\*\**p*=0.0068, F=7.38 by one-way ANOVA; **adjusted *p*=0.002) (G). Graph of co-localized CXCL5+/GLUT-1+ voxels in anti-IL-17B IgG-treated animals (*n=*4/grp; *\*p*=0.018). (H). Scale bars = 10 μm.

To block DIO-induced endothelial CXCL5 expression resulting from IL-17Rb activation, we employed repetitive peripheral injections of a function-blocking anti-IL-17B antibody or isotype control IgG for 6 weeks in Tie2-Cre;tdTomato mice on HFD (Fig. 4D, S1D). Endothelial CXCL5 expression within the tdT+ vasculature was reduced by 60.4% using this approach (Fig. 4H) while IL-17Rb levels were not changed (Fig. S5) indicating that DIO-induced increases in endothelial CXCL5 can be at least partially regulated through IL-17B signaling at the endothelial cell surface. Peripheral blocking of IL-17B signaling significantly reduced the association of OPCs with the cerebral vasculature in DIO mice (upper panels Fig. 4D, 4E, 4F). Vessel-associated OPC cell area was not significantly different in HFD mice administered anti-IL-17B antibody (lower panels Fig. 4D, Fig. 4G), suggesting additional HFD-associated pathways drive the regulation of OPC cell area.

To determine if downstream chemokine signaling pathways were active in OPC from DIO mice, we created a PDGFRα-CreERT:RiboTag conditional transgenic mouse. Administration of tamoxifen leads to robust expression of HA in PDGFRα+ OPCs and expression of HA remains in Olig2+ cells regardless of dietary condition (Fig. 5A). Fate mapping of PDGFRα+ OPCs one month after 12 weeks of DIO demonstrated no significant difference in the percentage of PDGFRα+/HA+ fate mapped cells (Fig. S7). In a separate cohort after induction of DIO, we pulsed with single dose tamoxifen and isolated white matter OPCs 48 hrs later by anti-HA pulldown followed by RNA-seq. In DIO HA+ OPCs, 1,932 genes were significantly up-regulated and 2,806 genes were down-regulated (FDR<0.05; Fig. 5B; SI Data File 3). Gene ontology of the differentially expressed genes (DEGs) induced in HA+ OPCs on HFD compared to the full murine genome enriched for multiple pathways involved in cell migration (Fig. 5C). Gene ontology of only DEGs revealed a role for OPCs in regulating endothelial cell proliferation in the setting of HFD-induced vascular changes (Fig. 5D). Pathway analysis demonstrated enrichment for downstream chemokine signaling with 31 of 198 chemokine signaling pathway genes (FDR = 1.62×10^-55^) (36) among the DEGs in HFD HA+ OPCs (Table 1).

**Figure 5.**
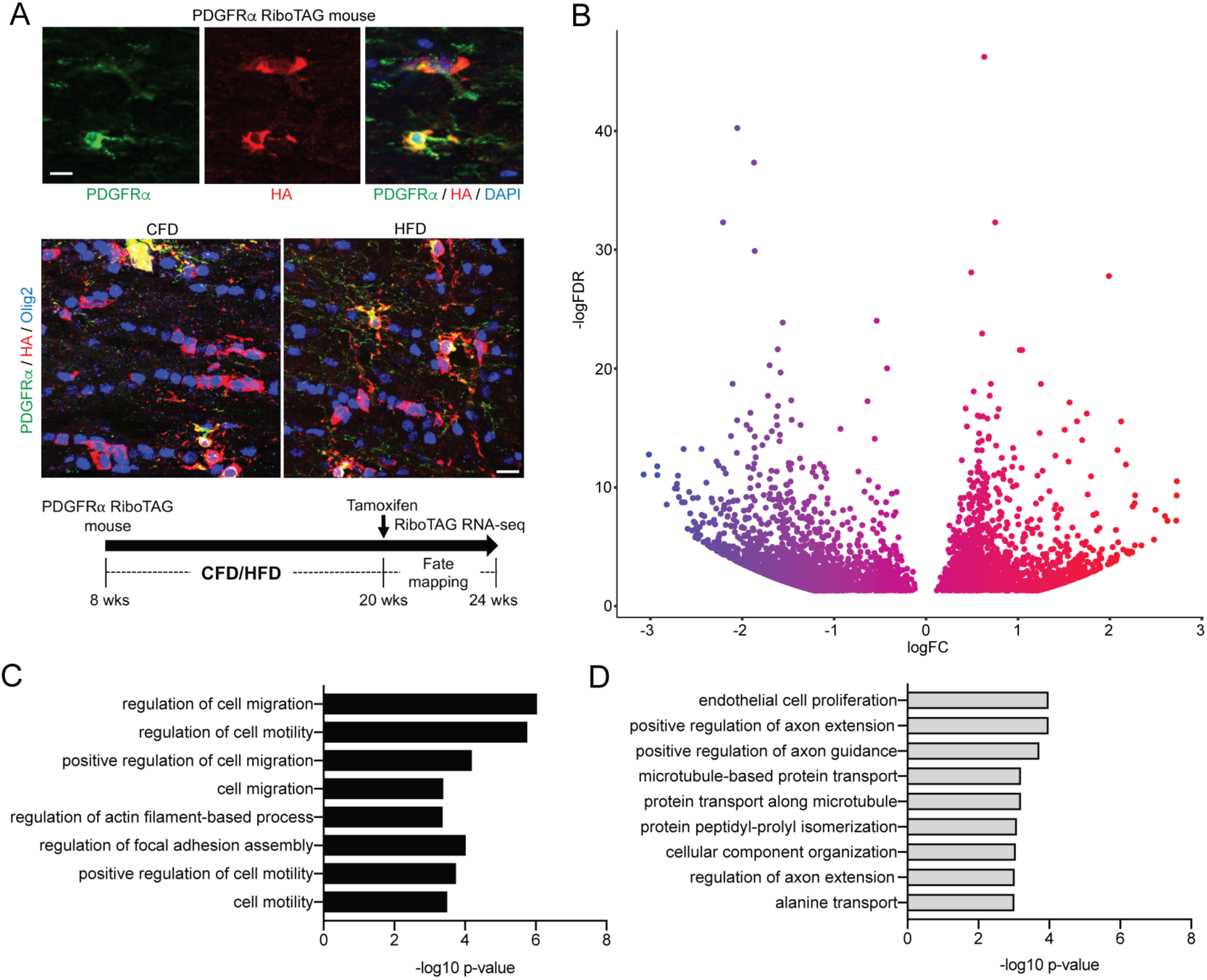
Transcriptional profiling of PDGFRα OPCs in DIO. Tamoxifen-induced expression of ribosome-associated hemagglutinin (HA; red) in PDGFRα+ OPCs (green) with restriction of HA expression to Olig2+ cells (blue) (A). Volcano plot of differentially expressed genes (FDR<0.05) induced by DIO in HA+ OPCs from PDGFRα RiboTAG mice (B). Enrichment values of gene ontology terms involved in cellular migration from the HFD OPC differentially expressed gene list (C). Top gene ontology terms enriched in PDGFRα+ OPCs after DIO (D). Complete gene list and ontology analysis available in SI Data File 3.

**Table 1.** Chemokine signaling pathway-related genes differentially expressed in DIO PDGFRα:RiboTAG OPCs

| Gene | logFC | adj. *p*-value | Location in chemokine signaling pathway |
| --- | --- | --- | --- |
| ADCY2 | 0.3783 | 0.00007 | cAMP signaling pathway |
| ADCY5 | 0.4331 | 0.03644 | cAMP signaling pathway |
| PRKACB | 0.2698 | 0.00240 | cAMP signaling pathway |
| CX3CL1 | 0.6773 | 0.00000 | Cytokine-cytokine receptor interaction |
| GRK3 | -0.2817 | 0.04388 | Cytokine-cytokine receptor interaction |
| GRK5 | -0.2740 | 0.01094 | Cytokine-cytokine receptor interaction |
| CCL27A | -1.1741 | 0.00916 | Cytokine-cytokine receptor interaction |
| CCR5 | 1.6514 | 0.00055 | Cytokine-cytokine receptor interaction |
| CCL25 | -1.0643 | 0.04174 | Cytokine-cytokine receptor interaction |
| CXCL12 | -1.3069 | 0.02963 | Cytokine-cytokine receptor interaction |
| RASGRP2 | 1.1673 | 0.01699 | Diacylglycerol pathway |
| GNG2 | 0.3214 | 0.04134 | Diacylglycerol pathway |
| GNB1 | -1.4145 | 0.00731 | Diacylglycerol pathway |
| GNB4 | -1.9793 | 0.00019 | Diacylglycerol pathway |
| GNB5 | -1.3031 | 0.02824 | Diacylglycerol pathway |
| STAT3 | -1.7856 | 0.00147 | Jak-STAT signaling pathway |
| SHC2 | -0.3250 | 0.03276 | MAPK signaling pathway |
| NRAS | -0.4424 | 0.00585 | MAPK signaling pathway |
| MAP2K1 | 0.2114 | 0.01887 | MAPK signaling pathway |
| SOS1 | 0.5083 | 0.00089 | MAPK signaling pathway |
| GSK3B | -1.5297 | 0.00350 | PIP-Akt signaling pathway |
| FOXO3 | 0.2776 | 0.01386 | PIP-Akt signaling pathway |
| PIK3CG | 1.3907 | 0.00916 | PIP-Akt signaling pathway |
| PIK3R5 | 1.3333 | 0.00095 | PIP-Akt signaling pathway |
| ROCK1 | -1.2784 | 0.03343 | Regulation of actin cytoskeleton |
| WASL | -1.8148 | 0.00012 | Regulation of actin cytoskeleton |
| RAC2 | 1.6040 | 0.00049 | Regulation of actin cytoskeleton |
| PARD3 | 0.4322 | 0.03699 | Regulation of actin cytoskeleton |
| ELMO1 | -1.4219 | 0.00125 | Regulation of actin cytoskeleton |
| PTK2 | -0.9568 | 0.04945 | Regulation of actin cytoskeleton/Diacylglycerol pathway |
| CRK | 0.4503 | 0.03329 | Regulation of actin cytoskeleton/Diacylglycerol pathway |

Differentially expressed genes (FDR<0.05) from HFD OPCs were filtered for those found in the KEGG Chemokine Signaling Pathway (KEGG Map #04062). Functional enrichment for chemokine signaling was established using a STRING database search (FDR for pathway enrichment = $1.62 \times 10^{-55}$ ). Gene name, log fold change (logFC), adjusted *p*-value (adj. *p*-value), and location of the gene within the signaling pathway are shown.

### Endothelial CXCL5 exaggerates the OPC response to focal white matter ischemia and impedes remyelination after stroke

To determine the consequence of endothelial CXCL5 on injury response after focal white matter ischemia, we used an established model of white matter stroke (37, 38) (Fig. 6A) that produces a distinct population of stroke-responsive PDGFRα+ OPCs (39). At 7d after white matter stroke, there was no significant difference in the stroke lesion volume comparing animals on CFD vs. HFD (Fig. S8). To determine the role that DIO-induced endothelial CXCL5 expression influences injury response after focal white matter ischemia, we labeled for GLUT-1 and CXCL5 at 7d post-stroke. We measured the percentage of CXCL5+ voxels that co-localized with GLUT-1 within the peri-infarct tissue surrounding the stroke. As in uninjured white matter, the percentage of CXCL5+/GLUT-1+ voxels were significantly increased within the peri-infarct tissue in animals on HFD (Fig. 6B). Immunofluorescent labeling for PDGFRα+ OPCs identified an increase in stroke-responsive OPCs per lesion in DIO mice compared to control (Fig. 6C) at 7d post-stroke. Spatial mapping of stroke-responsive OPCs coupled with nearest neighbor comparative analysis (Fig. S8) indicates a greater distribution of stroke-responsive OPCs that specifically occurs at the lesion margins in DIO mice compared to control (Fig. 6D) where endothelial CXCL5 levels were increased. This increase is at least partially accounted for by an increased association of OPCs with CXCL5+ vessel segments within the peri-lesional tissue. This DIO-induced OPC-vessel interaction in the early phase impacts repair after stroke. At 28d after stroke, we compared PDGFRα+ OPC and GST-π+ mature oligodendrocyte cell counts in three regions of interest spanning the ischemic white matter lesion. This analysis revealed a significant change in oligodendrocyte cell populations 28d after stroke (*p*=0.0011, two-way ANOVA, *F*=14.47) (Fig. 6E). An increased number of residual stroke-responsive PDGFRα+ OPCs were present at 28d post-stroke in animals on HFD compared to CFD (adjusted *p*=0.0114). The number of GST-π+ mature oligodendrocytes within the lesion at 28d post-stroke was variable and generally reduced in animals on HFD compared to CFD (adjusted *p*=0.0654).

**Figure 6.**
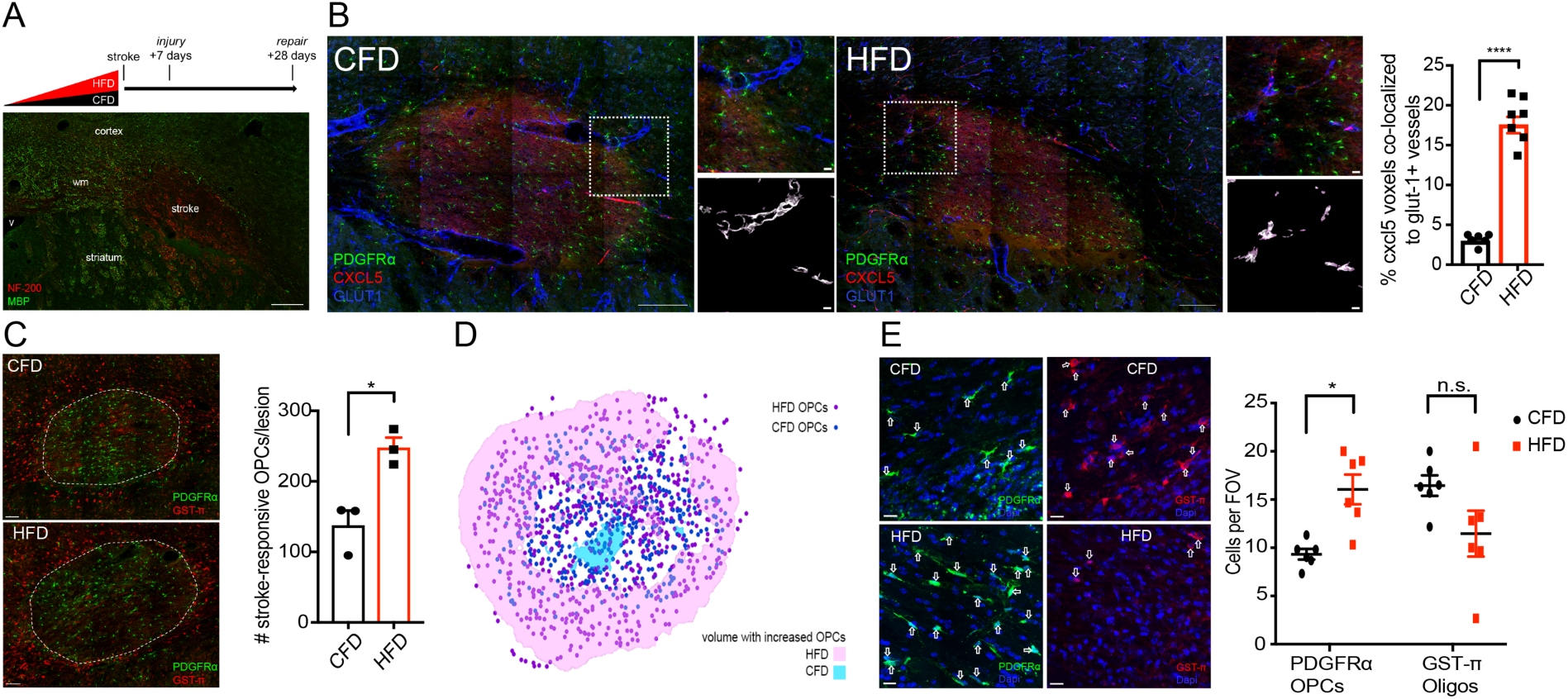
DIO-induced changes in stroke-responsive OPCs and repair after focal white matter stroke. Schematic of stroke modeling in animals on CFD and HFD (upper panel) with a representative white matter ischemic lesion shown with labeling for myelin basic protein (MBP, green) and neurofilament-200 (NF-200, red) (A). Labeling for GLUT-1 (blue), CXCL5 (red), and PDGFRα (green) at 7d post-stroke in animals on CFD (left) and HFD (right). Insets boxes from the peri-infarct tissue (upper) masked for GLUT-1 (white) with only co-localized CXCL5 (purple) (lower). Graph of percentage of co-localized CXCL5+/GLUT-1+ voxels (\*\*\*\**p*<0.0001) (B). Labeling of stroke-responsive PDGFRα+ OPCs (green) and GST-π (red) mature oligodendrocytes at 7d post-stroke and graph of #OPCs/lesion (C). Spatial mapping of stroke-responsive OPCs in CFD (dark blue) and HFD (purple) with shaded areas indicated regions of stroke lesion with statistically increased stroke-responsive OPCs between CFD (light blue) and HFD (pink) (D). Representative images from stroke-lesions at 28d post-stroke labeled for PDGFRα+ OPCs (green, left) and GST-π mature oligodendrocytes (red, right) between CFD (upper) and HFD (lower) animals with DAPI labeling of cell nuclei (blue). Graph of oligodendrocyte cell numbers at 28d post-stroke (\**p*=0.0114) (E). Scale bars = 10 μm. CFD and HFD.

### IL-17B and CXCL5 levels in human subjects at risk for cerebrovascular disease

Because we had identified that stimulation with IL-17B could drive endothelial secretion of CXCL5, we explored whether changes in circulating levels of IL-17B/IL-17Rb/CXCL5 signaling could be detected in human serum samples from a population at risk for small vessel disease. We utilized serum samples from the ASPIRE study cohort (40). We measured IL-17B and CXCL5 levels using a multiplexed Luminex assay in serum samples obtained at the time of their acute presentation after the development of neurologic symptoms (Fig. 7A). In those subjects with concurrent blood samples and MRI scans (*n*=131), subjects with detectable levels of IL-17B (*n*=32, mean IL-17B = 47.83 pg/mL) had higher median CXCL5 levels (1043.0 pg/mL) than in those without detectable IL-17B (*n=*99, 515.3 pg/mL; *p*<0.0001). In subjects with tissue-confirmed acute microvessel ischemic lesions, CXCL5 values were higher in those subjects with detectable IL-17B compared to those without measurable IL-17B levels (*p=*0.0157) (Fig. 7B). Median values for white matter hyperintensities measured by modified Fazekas scoring were higher in IL-17B+ subjects (3.0 vs. 2.0) though not statistically different (*p*=0.42) (Fig. 7C). BMI data was recorded on 81.6% of ASPIRE subjects and was not significantly different between IL-17B+ (26.5 kg/m^2^) and IL-17B-(25.6 kg/m^2^) subjects (*p*=0.53).

**Figure 7.**
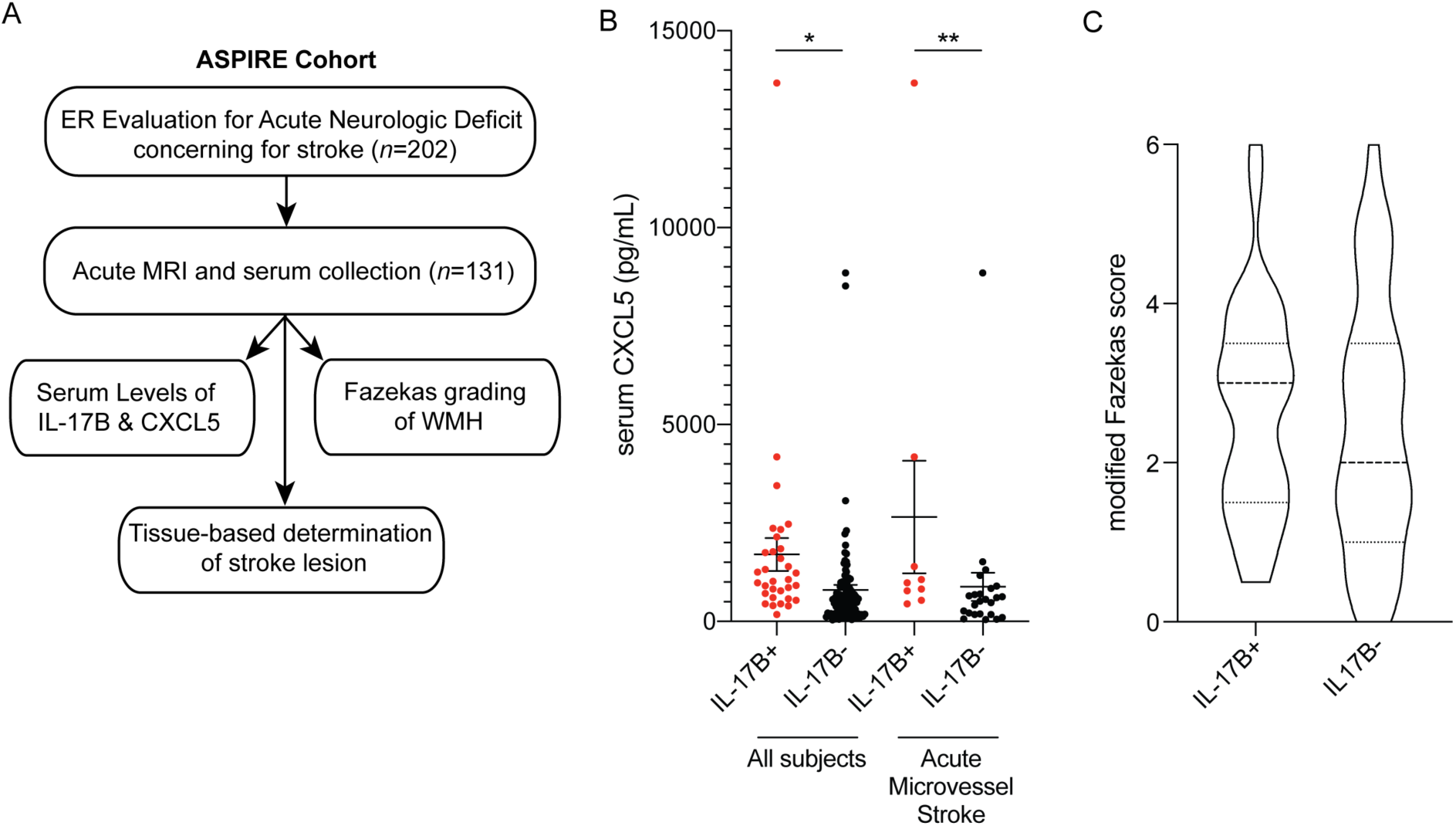
IL-17B/CXCL5 serum levels in subjects at risk for cerebrovascular disease. Subjects presenting for evaluation of acute neurologic deficits were enrolled in the ASPIRE study cohort (A). In all subjects, serum levels of CXCL5 were elevated in those subjects with detectable serum IL-17B (*n*=32; median 1043.0 pg/mL) compared to those with undetectable serum IL-17B (*n*=99; median 515.3 pg/mL; * *p*<0.0001; Mann-Whitney). Among those subjects with MRI-confirmed acute microvascular ischemia, serum CXCL5 levels were higher in IL-17B+ subjects (*n*=9; 978.2 pg/mL) vs. IL-17B-subjects (*n*=24; 539.7 pg/mL) (\*\**p*=0.0157; Mann Whitney) (B). Box and violin plot of modified Fazekas score in IL-17B+ vs. IL-17B-subjects (dashed line = median) (*p*=0.42; Mann-Whitney U-test) (C).

## Discussion

Cerebral small vessel disease is increasingly recognized as a substantial contributor to stroke risk and dementia (41). Microvascular injury in the brain is driven by cardiovascular risk factors yet molecular factors that link systemic vascular risk factors with molecular pathways in the brain are lacking. Obesity is a major cardiovascular and cerebrovascular risk factor, is growing in prevalence (42), is associated with white matter changes in humans (11, 13–15), and has a reliable animal model (27). Using this diet-induced obesity model, we show that obesity reduces white matter vasculature and increases OPCs in chronically injured white matter, as reported in other models (25). Our results showing ultrastructural changes in myelin in adult onset diet-induced obesity are similar to those seen in genetically obese (ob/ob) mice with reductions in myelin (43) and increases in OPCs in leptin-deficient ob/ob mice (44) validating this model for the study of chronic white matter injury. The present findings are the first to identify the transcriptome of chronically injured cerebral endothelia. We used that dataset to identify disordered vascular signaling that acts to regulate OPCs and impairs remyelination after stroke while also identifying a new signaling pathway relevant to human cerebral small vessel disease.

Cell-specific transcriptional profiling using ribosomal tagging is a valuable tool in parsing out molecular signals from a complex tissue such as the brain (31, 45). Here, we developed a methodology to profile cerebral endothelial cells *in vivo*. By combining this vascular Ribotag mouse with a chronic vascular risk factor model, we identified novel endothelial pathways that appear relevant to human cerebral small vessel disease. A similar vascular profiling approach could be easily applied to identify microvascular injury signals in other organs such as the kidney or retina. It could also be applied to other chronic or acute neurologic conditions that feature microvascular injury including aging, diabetes, or isolated hypertension. Here, we primarily focused on the paracrine signaling of chronically injured endothelial cells into the white matter and the consequence of this intercellular paracrine action on white matter repair. However, we also demonstrate that this vascular profiling approach may also facilitate the development of novel fluid-based biomarkers to track the response of the cerebral endothelium to chronic risk factors. Such efforts to better characterize the molecular pathways relevant to human cerebral small vessel disease is crucial for the development of diagnostic and therapeutic interventions (26).

Vessels and OPCs are known to interact both during development and to maintain white matter homeostasis (46). During CNS development, OPCs migrate extensively to distribute throughout the entire CNS and this migration requires the physical vascular scaffold (22). Cerebral endothelial cells secrete trophic factors that activate Src and Akt signaling pathways to support the survival and proliferation of OPCs (18). However, the full spectrum of molecular pathways that drive the vessel-OPC interaction remain largely unknown. The present data in disease and studies in the developing brain indicate that chemokines are critical. In-vivo time lapse imaging reveals that in the developing mouse brain, OPCs interact with vasculature and migrate along the vessels to the destined cerebral regions dependent on CXCR4 activation in OPCs, which binds to endothelial secreted ligand CXCL12, and promotes their attraction to cerebral vasculature (47). Our study illustrates a similar phenomenon, with DIO-induced endothelial expression of CXCL5 promoting the association of OPCs to the vasculature within adult white matter *in vivo*. Transcriptional profiling of OPCs in DIO using PDGFRα RiboTAG mice further imply that chemokine signaling pathways acting through CXCR2, the receptor for CXCL5, play a significant role in regulating a migratory interaction between endothelial cells (ECs) and OPCs. Based on gene ontology analysis, this interaction may promote angiogenesis and is critical to white matter homeostasis in the setting of chronic vascular injury.

What is the consequence of sequestering OPCs to the vascular bed? In the spontaneously hypertensive rat model of cerebral small vessel disease in which endothelial cells are progressively injured, OPCs are increased and fail to mature properly. In this model, dysfunctional endothelial cells impair the maturation of OPCs *in vitro* and promote their proliferation through the production of HSP90α (25). In the DIO mouse model, we observed a reduction in white matter microvascular complexity with the surviving endothelial cells responding with a specific transcriptional response implicating growth factor and immune signaling. In part, this response appears to cause OPCs to respond to vascular injury through CXCL5 signaling to OPCs. These OPCs are elongated along vessels with a long leading edge or display hypertrophied cell bodies and processes compared to those OPCs that are not associated with blood vessels, similar to those previously reported for migratory or reactive OPCs (22, 23, 50). These migratory OPCs are likely responding to disrupted vascular integrity and may facilitate endothelial cell proliferation but as a result of their new restriction to the vascular bed, fail to properly differentiate, ultimately compromising remyelination after stroke. Deactivation of chemokine signaling in OPCs has been shown to promote repair after autoimmune-mediated demyelination (48, 49). This disordered vascular regulation of myelination provides a new concept in understanding cellular signaling in cerebral small vessel disease. Though we clearly demonstrate that CXCL5 can serve as this vascular injury regulatory signal in rodents and humans, further studies are needed to definitively prove whether this response serves some partially protective role on the blood-brain barrier through reciprocal OPC to endothelial signaling (51).

White matter ischemic lesions are characterized by a robust early loss of axons, myelin and oligodendrocytes (52–54). Similar to inflammatory white matter lesions (55), OPCs respond early and robustly to white matter ischemic lesions common to the aging human brain (39, 53, 56). The peri-infarct white matter at the margin of the ischemic lesion, often referred to as the white matter penumbral region (57), is where cross-talk between axons and oligodendrocytes is compromised (58, 59). Here, we used a novel approach to identify the spatial relationship of stroke-responsive OPCs to a focal white matter stroke lesion. This approach directly informs our data by showing that the increase in stroke-responsive OPCs produced by obesity occurs precisely in the peripheral margins of the white matter stroke lesion where tissue repair and the stimuli for remyelination would be maximal. In DIO mice, the stroke-responsive OPC lesion area is 30% larger and this expanded penumbral region is marked by increased endothelial CXCL5 expression explaining why more stroke-responsive OPCs are seen at the lesion periphery. The consequence of this is apparent at 28 days after stroke; DIO-induced endothelial expression of CXCL5 leaves behind a population of activated, injury-responsive OPCs whose maturation is inhibited. This progenitor restricted state could indicate that remyelination is simply delayed after stroke or it could lead to a progressive dysfunctional OPC response as the ability of NG2+ OPCs to differentiate into oligodendrocytes declines with chronic insults (60).

From our data, an emerging concept places the cerebral endothelial cell at the center of the pathophysiology relevant to cerebral small vessel disease. As the conduit between the brain and systemic insults such as hypertension, diabetes, and the metabolic disturbances of obesity, identifying molecular pathways in the cerebral endothelia represent an attractive target for understanding disease pathogenesis. Changes in normal white matter homeostasis that result from chronic cerebrovascular risk factors can directly alter injury response and repair after stroke by acting through vascular regulation of myelination. That the pathway we characterized is functionally absent from normal young adult mice yet ubiquitous in aged human brain and detectable in the serum of subjects at risk for cerebral small vessel disease suggests greater efforts to appropriately model co-morbid conditions in animal models of stroke and cerebrovascular injury may pave a smoother path to translation into human trials.

## Material and Methods

### Animals

Mice were housed under UCLA regulation with a 12-hour dark-light cycle. All animal studies presented here were approved by the UCLA Animal Research Committee ARC#2014-067-01B, accredited by the AAALAC. Diet-induced obesity was induced in mice by ad lib feeding with 60%kCal from fat chow (HFD) or 10%kCal from fat chow (CFD) (Research Diets, Inc.). Weights (g) were measured weekly. Mice strains used in this study are described in *SI Materials and Methods*.

### Translating ribosome affinity purification and RNA-sequencing

HA-tagged ribosomal associated RNAs from cerebral white matter endothelia or OPCs were isolated and purified by Nucleospin miRNA kit (Machary-Nagel). RNA-sequencing was run using 69 bp paired end reads. Reads were aligned to the mouse genome using STAR (v.mm10). Differential gene expression analysis was performed using EdgeR assuming an FDR <0.1 as significant. Gene ontology analysis was performed using GOrilla (61) and Enrichr (62). Additional information of RNA isolation, RNA sequencing and analysis are described in *SI Materials and Methods*.

### Human brain microvascular endothelial cell culture

Primary Human Brain Microvascular Endothelial Cells (HBMECs) (Cell Systems) between P5-P9 were maintained at 37°C until confluence with manufacturer recommended media containing serum with media exchange every two days. Maintenance cultures were re-plated into a 96-well filter bottom plate and cultured until near confluence. Two days after seeding, HBMECs were stimulated with culture medium containing 250 ng/ml of mouse IL-17A, B, C, D, or E (R&D Systems, Inc.). Conditioned media from triplicate culture conditions was collected after 48 hours and human CXCL5 levels measured using a human CXCL5 Quantikine Elisa Kit (R&D Systems, Inc.).

### Immunofluorescence and confocal imaging

Animals were euthanized with a lethal dose of isoflurane, transcardially perfused with PBS followed by 4% paraformaldehyde in 0.1 M sodium phosphate buffer, brains removed, post-fixed for 24 hrs and cryoprotected for 48 hrs in 30% sucrose in PBS. Forty microns coronal cryosections and immunostaining was performed essentially as described (37). Details regarding antibodies and microscopic imaging are available in *SI Materials and Methods*. Human post-mortem brain sections were selected from the UC Davis ADC Neuropathology Core samples based on *a priori* selection criteria and stained for CXCL5/6 using standard immunohistochemistry.

### White matter stroke

Subcortical white matter ischemic injury was induced as previously described (38) using three stereotactic injections of L-Nio (L-N^5^-(1-Iminoethyl) ornithine, dihydrochloride; Calbiochem) into the subcortical white matter under sensorimotor cortex. Animals (*n* = 4/grp) were sacrificed at 7- or 28-days post-stroke and analyzed for tissue outcomes.

### Lentiviral injection

A dual promoter lentiviral backbone was created through sequential subcloning to place either GFP or murine CXCL5 between loxP sites. Control GFP and CXCL5-GFP lentivirus were packaged in human 293 cells (ATCC cat. no. CRL-11268) and concentrated by ultracentrifugation on a sucrose column. 200 nL of concentrated virus was injected into the subcortical white matter and allowed to express for 6 weeks.

### Anti-IL-17B antibody administration

Anti-mIL-17B function blocking antibody (R&D, AF1709) was diluted with 0.9% saline to a concentration of 1mg/ml. Normal Goat isotype-matched IgG (R&D, AB-108-C) was used as control. Tie2-Cre;tdTomato mice were fed with high fat diet starting at 8 weeks old and weighed weekly. Aliquots of 50μg of anti-mIL-17B IgG or control IgG were prepared and administered in a blinded fashion every 72 hours by intraperitoneal injection from 14 weeks old and analyzed 48 hours after the last injection at 20 weeks old.

### ASPIRE Study Cohort

Research involving human subjects was approved by the UCLA Institutional Review Board (IRB # 14-001798) and was conducted in compliance with the Health Information Portability and Accountability Act. Serum levels of IL-17B and CXCL5 were measured in duplicate using a custom assay on the Luminex platform (R&D Systems). The manufacturer protocol was followed and antigen binding within the assay was measured on a Luminex 200 System and analyzed using Milliplex Analyst 5.1. Modified Fazekas scores were determined by blinded analysis of T_2_-weighted FLAIR images.

### Statistical analysis

The number of animals used in each experiment is listed in the Results section or Figure Legend. Microscopic tissue analysis was performed in a blinded fashion. Statistical analysis was performed using GraphPad Prism 7 software. Unless otherwise stated, statistical significance was determined using α=0.05 using non-parametric approaches when applicable and corrected for multiple comparisons. Data are shown as mean ± SEM.

### Other Methods

Further details regarding procedures related to gene expression and analysis, immunohistochemistry, electronic microscopy, spatial analysis, human specimen and imaging analysis, and statistical analysis are explained in detail in *SI Materials and Methods*. All DNA sequences, primers, plasmids and packaged viruses are available upon request. Gene expression data is available in the *SI Data Files.* ASPIRE study data is available upon request.

## Author contributions

G.X., Ro.K., and J.D.H. designed research and collected data; G.X., Ri.K., Y.K., and J.D.H. performed gene expression analysis; G.X. and J.B. performed spatial analysis; G.X., L.A.H., and J.D.H. performed electron microscopy and analysis; A.B., M.M, S.T.C., and J.D.H performed lentiviral cloning and packaging; P.M, N.K. and J.D.H. performed *in vitro* cell culture experiments and analysis; C.K.W, X.R.Z, H.V.V., V.K, G.X, and C.D. performed human brain tissue selection, staining, and analysis; V.K., I.A., S.A.S., and J.D.H. enrolled subjects, extracted data, and performed imaging and biospecimen analysis; G.X. and J.D.H. wrote the paper.

## Acknowledgments

The authors are grateful to Kelsey Ericson for preparation of human brain tissue specimens and the members of the UCLA Stroke Force and Stroke Center members who assisted in ASPIRE Trial enrollment and data collection. This work was graciously supported by grants from the Larry L. Hillblom Foundation (TLLHF 2014-A-014) and the American Heart Association (#15CRP22900006; 16GRNT31080021). LAH and STC receive support from the Dr. Miriam and Sheldon G. Adelson Medical Research Foundation. CD receives support from UC Davis Alzheimer’s Disease Center grant P30 AG 010129. JDH received support from the National Institute for Neurological Disorders and Stroke (K08 NS083740) and the United States Department of Veterans Affairs Greater Los Angeles Healthcare System.

## References

1. Gouw AA, et al. (2008) Progression of white matter hyperintensities and incidence of new lacunes over a 3-year period: the Leukoaraiosis and Disability study. Stroke 39(5):1414–1420.

2. Inzitari D, et al. (2009) Changes in white matter as determinant of global functional decline in older independent outpatients: three year follow-up of LADIS (leukoaraiosis and disability) study cohort. BMJ 339:b2477.

3. Jokinen H, et al. (2011) Incident lacunes influence cognitive decline: the LADIS study. Neurology 76(22):1872–1878.

4. Koga H, et al. (2009) Cognitive consequences of multiple lacunes and leukoaraiosis as vascular cognitive impairment in community-dwelling elderly individuals. J Stroke Cerebrovasc Dis 18(1):32–37.

5. Reed BR, et al. (2004) Effects of white matter lesions and lacunes on cortical function. Arch Neurol 61(10):1545–1550.

6. Debette S & Markus HS (2010) The clinical importance of white matter hyperintensities on brain magnetic resonance imaging: systematic review and meta-analysis. BMJ 341:c3666.

7. van Dijk EJ, et al. (2004) The association between blood pressure, hypertension, and cerebral white matter lesions: cardiovascular determinants of dementia study. Hypertension 44(5):625–630.

8. de Leeuw FE, et al. (2002) Hypertension and cerebral white matter lesions in a prospective cohort study. Brain 125(Pt 4):765–772.

9. Jongen C, et al. (2007) Automated measurement of brain and white matter lesion volume in type 2 diabetes mellitus. Diabetologia 50(7):1509–1516.

10. Manschot SM, et al. (2006) Brain magnetic resonance imaging correlates of impaired cognition in patients with type 2 diabetes. Diabetes 55(4):1106–1113.

11. Bokura H, Yamaguchi S, Iijima K, Nagai A, & Oguro H (2008) Metabolic syndrome is associated with silent ischemic brain lesions. Stroke 39(5):1607–1609.

12. Park K & Yasuda N (2009) Association between metabolic syndrome and minimal leukoaraiosis. Stroke 40(1):e5; author reply e6.

13. Park K, et al. (2008) Significant associations of metabolic syndrome and its components with silent lacunar infarction in middle aged subjects. J Neurol Neurosurg Psychiatry 79(6):719–721.

14. Yin ZG, et al. (2014) Association between metabolic syndrome and white matter lesions in middle-aged and elderly patients. Eur J Neurol 21(7):1032–1039.

15. Kwon HM, et al. (2009) Significant association of metabolic syndrome with silent brain infarction in elderly people. J Neurol 256(11):1825–1831.

16. Fisher CM (1979) Capsular infarcts: the underlying vascular lesions. Arch Neurol 36(2):65–73.

17. Esiri MM, Wilcock GK, & Morris JH (1997) Neuropathological assessment of the lesions of significance in vascular dementia. J Neurol Neurosurg Psychiatry 63(6):749–753.

18. Arai K & Lo EH (2009) An oligovascular niche: cerebral endothelial cells promote the survival and proliferation of oligodendrocyte precursor cells. J Neurosci 29(14):4351–4355.

19. Pham LD, et al. (2012) Crosstalk between oligodendrocytes and cerebral endothelium contributes to vascular remodeling after white matter injury. Glia 60(6):875–881.

20. Maki T, et al. (2015) Potential interactions between pericytes and oligodendrocyte precursor cells in perivascular regions of cerebral white matter. Neurosci Lett 597:164–169.

21. Marques S, et al. (2016) Oligodendrocyte heterogeneity in the mouse juvenile and adult central nervous system. Science 352(6291):1326–1329.

22. Tsai HH, et al. (2016) Oligodendrocyte precursors migrate along vasculature in the developing nervous system. Science 351(6271):379–384.

23. Hayakawa K, et al. (2011) Vascular endothelial growth factor regulates the migration of oligodendrocyte precursor cells. J Neurosci 31(29):10666–10670.

24. Hayakawa K, et al. (2012) Cerebral endothelial derived vascular endothelial growth factor promotes the migration but not the proliferation of oligodendrocyte precursor cells in vitro. Neurosci Lett 513(1):42–46.

25. Rajani RM, et al. (2018) Reversal of endothelial dysfunction reduces white matter vulnerability in cerebral small vessel disease in rats. Sci Transl Med 10(448).

26. Corriveau RA, et al. (2017) Alzheimer’s Disease-Related Dementias Summit 2016: National research priorities. Neurology 89(23):2381–2391.

27. Surwit RS, Kuhn CM, Cochrane C, McCubbin JA, & Feinglos MN (1988) Diet-induced type II diabetes in C57BL/6J mice. Diabetes 37(9):1163–1167.

28. Kennedy AJ, Ellacott KL, King VL, & Hasty AH (2010) Mouse models of the metabolic syndrome. Dis Model Mech 3(3-4):156–166.

29. Huang PL (2009) A comprehensive definition for metabolic syndrome. Dis Model Mech 2(5-6):231–237.

30. Zhang Y, et al. (2014) An RNA-sequencing transcriptome and splicing database of glia, neurons, and vascular cells of the cerebral cortex. J Neurosci 34(36):11929–11947.

31. Sanz E, et al. (2009) Cell-type-specific isolation of ribosome-associated mRNA from complex tissues. Proc Natl Acad Sci U S A 106(33):13939–13944.

32. Gaffen SL (2009) Structure and signalling in the IL-17 receptor family. Nat Rev Immunol 9(8):556–567.

33. Chen K, et al. (2016) IL-17 Receptor Signaling in the Lung Epithelium Is Required for Mucosal Chemokine Gradients and Pulmonary Host Defense against K. pneumoniae. Cell Host Microbe 20(5):596–605.

34. Strieter RM, et al. (1995) The functional role of the ELR motif in CXC chemokine-mediated angiogenesis. J Biol Chem 270(45):27348–27357.

35. Wang LY, Tu YF, Lin YC, & Huang CC (2016) CXCL5 signaling is a shared pathway of neuroinflammation and blood-brain barrier injury contributing to white matter injury in the immature brain. J Neuroinflammation 13:6.

36. Szklarczyk D, et al. (2019) STRING v11: protein-protein association networks with increased coverage, supporting functional discovery in genome-wide experimental datasets. Nucleic Acids Res 47(D1):D607–D613.

37. Hinman JD, Rasband MN, & Carmichael ST (2013) Remodeling of the axon initial segment after focal cortical and white matter stroke. Stroke 44(1):182–189.

38. Nunez S, et al. (2016) A Versatile Murine Model of Subcortical White Matter Stroke for the Study of Axonal Degeneration and White Matter Neurobiology. J Vis Exp (109).

39. Sozmen EG, et al. (2016) Nogo receptor blockade overcomes remyelination failure after white matter stroke and stimulates functional recovery in aged mice. Proc Natl Acad Sci U S A 113(52):E8453–E8462.

40. Azizkhanian I, et al. (2019) Plasma Lipid Profiling Identifies Biomarkers of Cerebral Microvascular Disease. Front Neurol 10:950.

41. Debette S, Schilling S, Duperron MG, Larsson SC, & Markus HS (2018) Clinical Significance of Magnetic Resonance Imaging Markers of Vascular Brain Injury: A Systematic Review and Meta-analysis. JAMA Neurol.

42. Services USDoHH (2018) Nutrition, Physical Activity, and Obesity: Data, Trends and Maps.

43. Sena A, Sarlieve LL, & Rebel G (1985) Brain myelin of genetically obese mice. J Neurol Sci 68(2-3):233–243.

44. Udagawa J, Nimura M, & Otani H (2006) Leptin affects oligodendroglial development in the mouse embryonic cerebral cortex. Neuro Endocrinol Lett 27(1-2):177–182.

45. Heiman M, Kulicke R, Fenster RJ, Greengard P, & Heintz N (2014) Cell type-specific mRNA purification by translating ribosome affinity purification (TRAP). Nat Protoc 9(6):1282–1291.

46. Arai K & Lo EH (2009) Oligovascular signaling in white matter stroke. Biol Pharm Bull 32(10):1639–1644.

47. Banisadr G, et al. (2011) The role of CXCR4 signaling in the migration of transplanted oligodendrocyte progenitors into the cerebral white matter. Neurobiol Dis 44(1):19–27.

48. Marro BS, et al. (2019) Disrupted CXCR2 Signaling in Oligodendroglia Lineage Cells Enhances Myelin Repair in a Viral Model of Multiple Sclerosis. J Virol 93(18).

49. Liu L, et al. (2010) Myelin repair is accelerated by inactivating CXCR2 on nonhematopoietic cells. J Neurosci 30(27):9074–9083.

50. Hampton DW, Rhodes KE, Zhao C, Franklin RJ, & Fawcett JW (2004) The responses of oligodendrocyte precursor cells, astrocytes and microglia to a cortical stab injury, in the brain. Neuroscience 127(4):813–820.

51. Seo JH, et al. (2014) Oligodendrocyte precursor cells support blood-brain barrier integrity via TGF-beta signaling. PLoS One 9(7):e103174.

52. Back SA, et al. (2007) Hypoxia-ischemia preferentially triggers glutamate depletion from oligodendroglia and axons in perinatal cerebral white matter. J Cereb Blood Flow Metab 27(2):334–347.

53. Sozmen EG, Kolekar A, Havton LA, & Carmichael ST (2009) A white matter stroke model in the mouse: axonal damage, progenitor responses and MRI correlates. J Neurosci Methods 180(2):261–272.

54. Valeriani V, Dewar D, & McCulloch J (2000) Quantitative assessment of ischemic pathology in axons, oligodendrocytes, and neurons: attenuation of damage after transient ischemia. J Cereb Blood Flow Metab 20(5):765–771.

55. Tripathi RB, Rivers LE, Young KM, Jamen F, & Richardson WD (2010) NG2 glia generate new oligodendrocytes but few astrocytes in a murine experimental autoimmune encephalomyelitis model of demyelinating disease. J Neurosci 30(48):16383–16390.

56. Miyamoto N, et al. (2015) Astrocytes Promote Oligodendrogenesis after White Matter Damage via Brain-Derived Neurotrophic Factor. J Neurosci 35(41):14002–14008.

57. Maillard P, et al. (2011) White matter hyperintensity penumbra. Stroke 42(7):1917–1922.

58. Hinman JD, Lee MD, Tung S, Vinters HV, & Carmichael ST (2015) Molecular disorganization of axons adjacent to human lacunar infarcts. Brain 138(Pt 3):736–745.

59. Rosenzweig S & Carmichael ST (2015) The axon-glia unit in white matter stroke: mechanisms of damage and recovery. Brain Res 1623:123–134.

60. Mason JL, et al. (2004) Oligodendrocytes and progenitors become progressively depleted within chronically demyelinated lesions. Am J Pathol 164(5):1673–1682.

61. Eden E, Navon R, Steinfeld I, Lipson D, & Yakhini Z (2009) GOrilla: a tool for discovery and visualization of enriched GO terms in ranked gene lists. BMC Bioinformatics 10:48.

62. Gundersen GW, et al. (2015) GEO2Enrichr: browser extension and server app to extract gene sets from GEO and analyze them for biological functions. Bioinformatics 31(18):3060–3062.

