## Supplemental Information for "Endothelial CXCL5 negatively regulates myelination and repair after white matter stroke"

Ph: 310-825-6761

**Data and materials availability**: All DNA sequences, primers, plasmids and packaged viruses are available upon request. Gene expression data is available in the Supplemental Data Files.

S

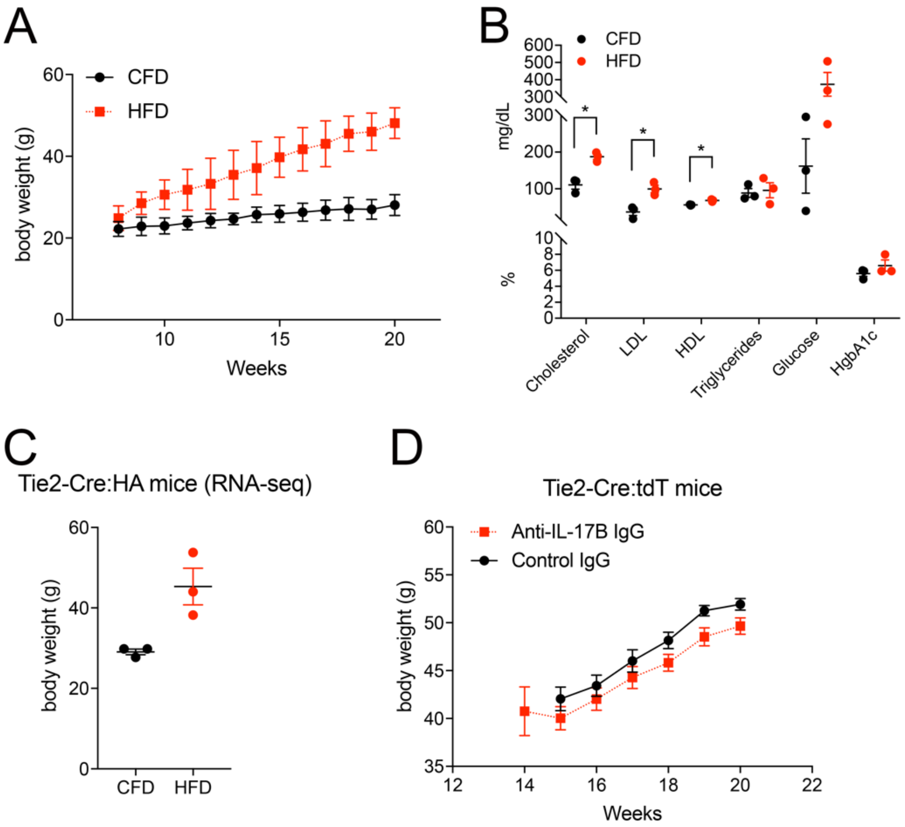


**Supplemental Figure 1. High fat diet induces obesity in mice.**
Week by week body weight measurements of C57BL/6J fed ad lib with CFD (black) or HFD (red) with final average 84.0% weight gain compared to 21.7% in CFD mice (***p*=0.0005; *n*=5) (A). Serologic testing of C57BL/6J mice on CFD or HFD (*n*=3) for total cholesterol (110.7±2.6 vs. 187.3±2.5; **p*=0.023), LDL (37.0 ±2.4 vs. 99.7±2.0; **p*=0.043), glucose (162.0±6.5 vs. 373.3±4.2) and HgbA1c (5.6 ±0.5vs. 6.6±0.2) (B). Final 20 week old body weights of Tie2-Cre:RiboTag HFD mice used for RNA-seq endothelial profiling (45.3±1.9 g vs. 29.1±0.6 g; *n*=3) (C). Final 20 week old body weights of Tie2-Cre;tdTomato mice fed ad lib on HFD treated with control IgG (black) and anti-IL-17B IgG (red) (51.9±0.5 g vs 49.1±0.6 g; **p*=0.029; *n*=4) (D). Data are mean ± SEM.

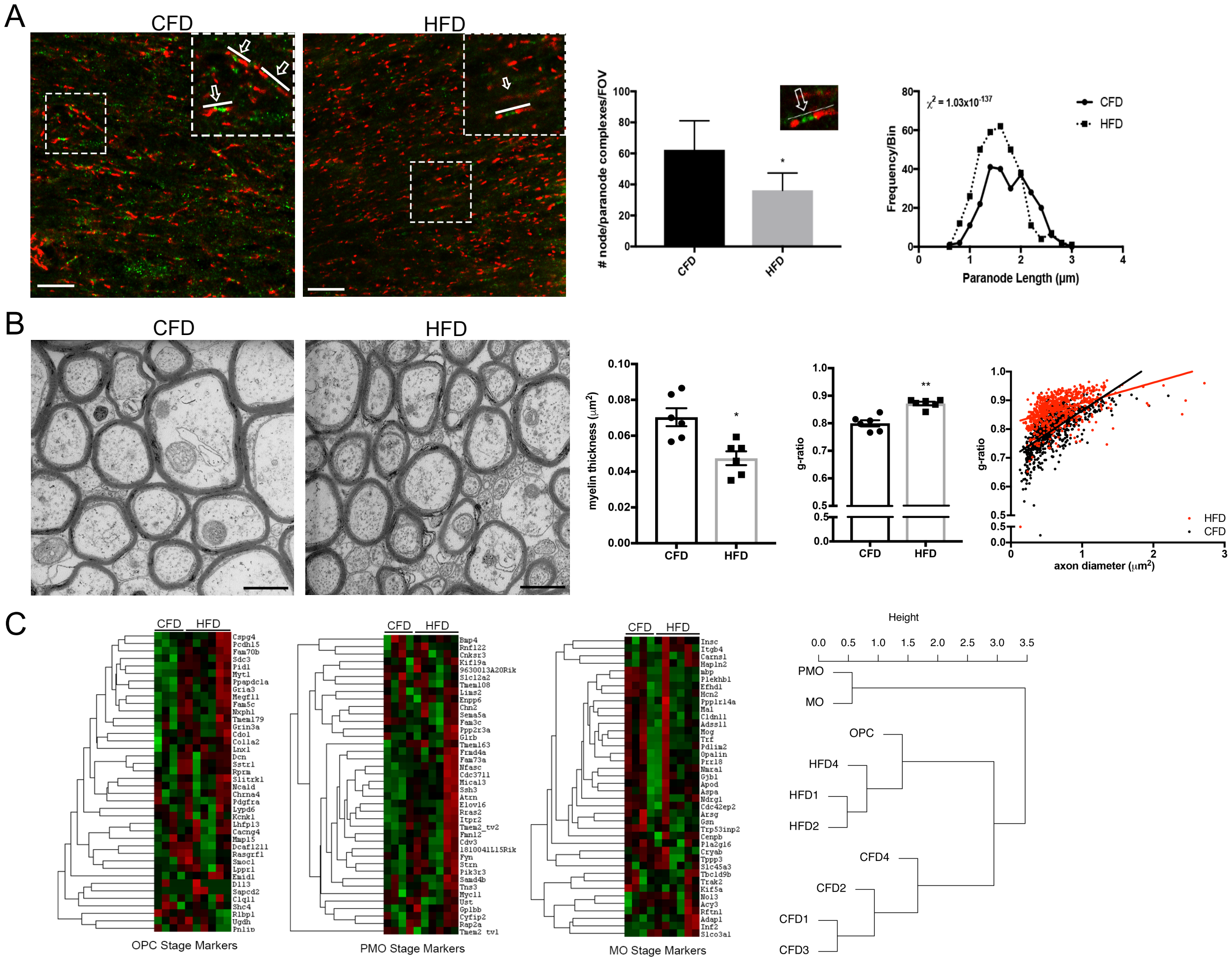


**Supplemental Figure 2. Validation of white matter disruption in HFD mice.**
Labeling for nodal ($Na_V$1.6, green) and paranodal (caspr, red) axonal microdomain segments within subcortical white matter in animals on CFD (left) and HFD (right). Inset boxes show examples of intact nodal and paranodal complexes (arrow with line). Average number of nodal and paranodal complexes, defined as adjacent caspr+ paranodal segments with concurrent $Na_V$1.6+ nodal segments (example in inset) (62.3±13.3 vs 36.2±7.9, ***$p$=1.7666 x $10^{-5}$; $n$=2). Distribution of paranodal length (binned by 0.2 µm) in animals on HFD (dashed line) compared to CFD (solid line) demonstrating a significant left shift towards shorter paranodes in animals on HFD (1.29±0.02 µm vs 1.70±0.03 µm, Chi-square=1.7117 x $10^{-138}$) (A). Callosal fibers at the midline were prepared in sagittal section for electron microscopy. Animals on CFD ($n$=6) demonstrated regular axonal and fiber diameter with an abundance of normally myelinated fibers (left panel, B). After 12 weeks on HFD ($n$=6), myelin ultrastructure in callosal fibers is compromised with reductions in myelin sheath thickness without significant changes in axonal number or fiber diameter (right panel, B). Myelin sheath thickness was significantly reduced (0.070µm vs. 0.047µm; *$p$=0.0087) (B). The average g-ratio was increased (0.88 vs. 0.80; **$p$=0.002) in animals on HFD compared to CFD (B). The distribution of axon diameter vs. g-ratio for individual axon measurements in animals on CFD (black) vs. HFD (red) showing that axon diameter was not significantly different between the groups (0.62±0.06 µm vs 0.69±0.05 µm; $p$=0.48) (B). Heatmap of oligodendrocyte lineage stage (OPC = oligodendrocyte progenitor

cell; PMO = premyelinating oligodendrocyte; MO = myelinating oligodendrocyte) gene expression profiles generated by Nanostring Oligodendrocyte Staging Assay (total 120 genes/40 per stage) for animals on CFD and HFD (*n*=4) (C), upregulated genes (red), downregulated genes (green); Hierarchical clustering of samples compared to reference marker gene expression profiles (Zhang et al. *J. Neurosci* 2014) shows that gene expression profiles of animals on HFD cluster with OPCs (C). Data are mean ± SEM. Scale bars = 10 μm (A), 1 μm (7200X magnification) (B).

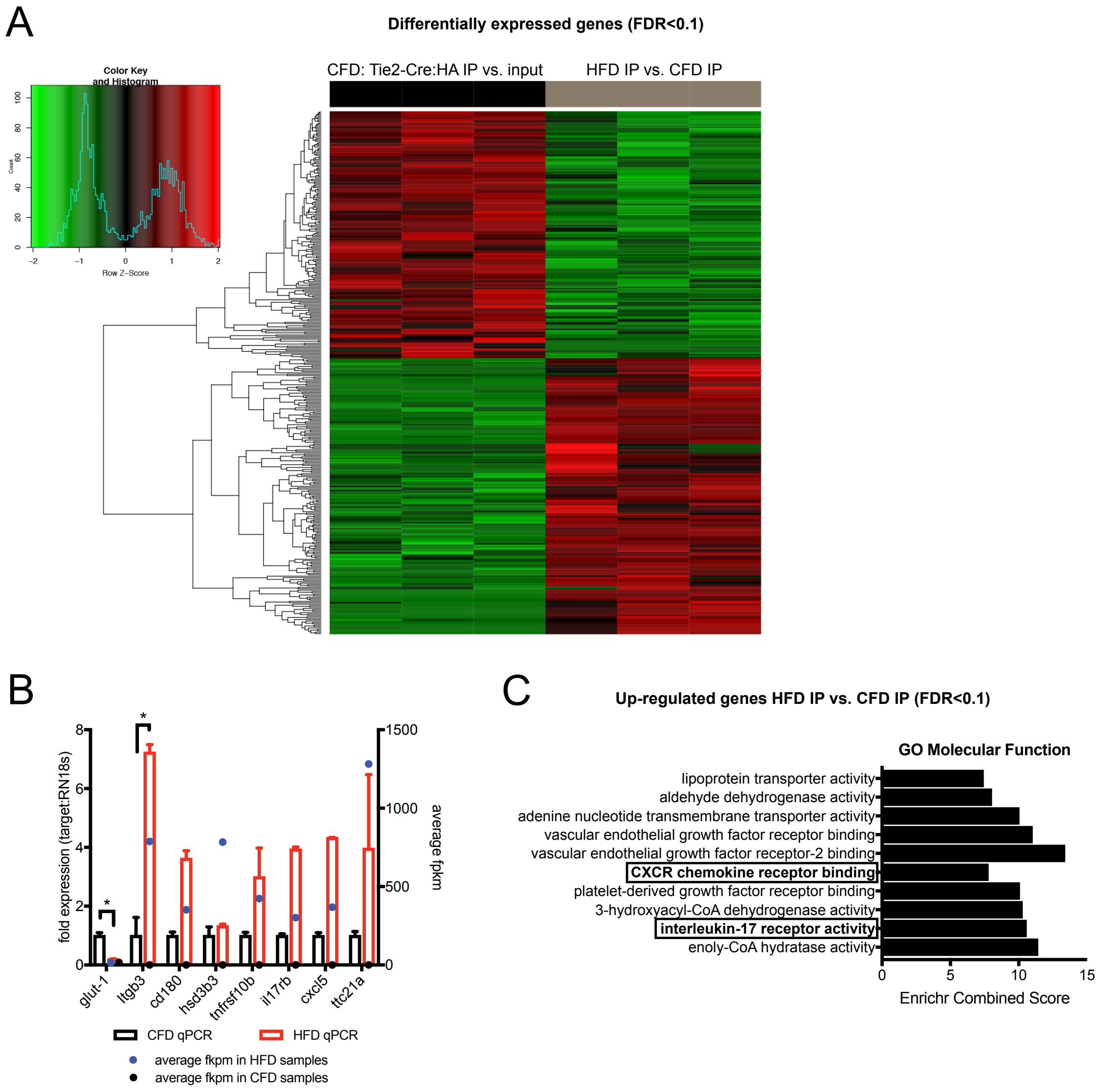


**Supplemental Figure 3. DIO results in a specific white matter endothelial gene expression profile and immune signaling activation.**

Heatmap representation of differentially expressed genes (FDR<0.1) between Tie2-Cre:RiboTag anti-HA immunoprecipitated ribosomes (IP) and total white matter (input) in animals on CFD (left, black) demonstrating unique transcriptional profile of white matter endothelia. Differentially expressed genes between Tie2-Cre:RiboTag anti-HA immunoprecipitated ribosomes (IP) on CFD vs. HFD (right, gray). Down-regulated genes shown in green and up-regulated genes shown in red (A). TRAP-qPCR confirmation of RNA-sequencing results for selected genes in independent Tie2-Cre:RiboTag mice ($n$=2/grp) (*$p$<0.05) (B). Bar plot of the Enrichr combined score of the gene ontology (molecular function) of up-regulated genes in Tie2-Cre:RiboTag anti-HA immunoprecipitated ribosomes (IP) HFD mice indicates enrichment of immune signaling pathways including C-X-C chemokine signaling and interleukin-17 receptor activation (C).

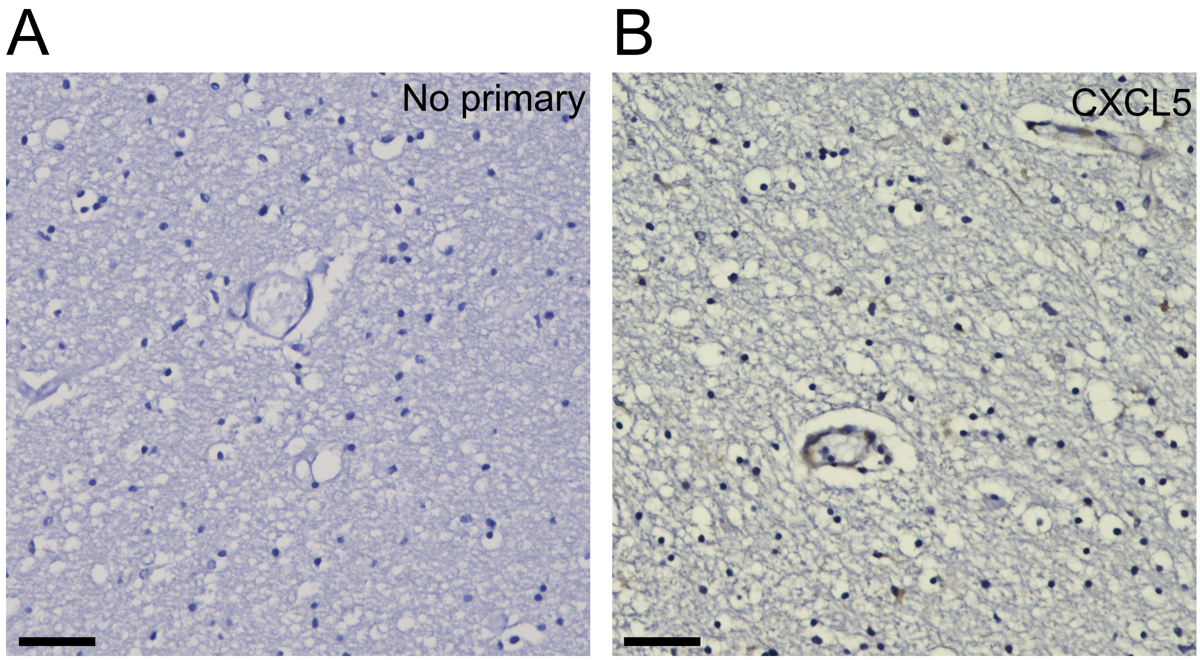


C

| Age | Sex | Ethnicity | Clinical Syndrome | Pathological Diagnosis | BRAAK | PMI | Any positive vessel | % positive vessel segments (# positive) |
|---|---|---|---|---|---|---|---|---|
| 75 | M | Hispanic | No cognitive impairment | Normal Brain | 2 | 49.5 | Yes | 35.3% (6) |
| 83 | M | White | No cognitive impairment | Possible AD (CERAD) | 2 | 30 | Yes | 60.0% (9) |
| 96 | F | White | No cognitive impairment | Normal Brain | 2 | 6.9 | Yes | 94.4% (17) |
| 86 | F | White | Questionable cognitive impairment | Vascular disease, infarcts only | 1 | - | No | 0% (0) |
| 84 | F | White | No cognitive impairment | Normal Brain | 1 | 10.9 | Yes | 100% (24) |
| 72 | M | White | No cognitive impairment | Vascular disease, combined lacunar and large infarctions, cerebrovascular disease | 1 | 4 | Yes | 50.0% (9) |
| 89 | M | Hispanic | Dementia | Vascular disease, multiple lacunes | 2 | 69 | Yes | 71.4% (15) |
| 92 | F | White | Dementia | Vascular disease, multiple lacunes | 3 | 7 | Yes | 50.0% (7) |
| 90 | F | Asian | Dementia | Vascular disease, Binswangers disease. Pathological findings of white matter vasculopathy | 1 | 40 | No | 0% (0) |
| 92 | F | Hispanic | Dementia | Limbic sclerosis, entorhinal neocortex | 3 | 31 | Yes | 84.6% (11) |

**Supplemental Figure 4. Detection of CXCL5 in human peri-ventricular white matter.**

Immunohistochemistry without anti-CXCL5 antibody demonstrates no visible staining in periventricular white matter (A). Immunohistochemistry for CXCL5 in periventricular white matter reveals clear detection of CXCL5 (brown) within endothelial cells in the white matter (B). CXCL5 staining is also apparent in scattered glial cells within the white matter. Scale bars = 50 µm. Cases selected from a convenience cohort (*n*=10) of a large database of post-mortem subjects from the University of California Davis ADRC. These subjects were selected to include those with and without cognitive impairment (clinical syndrome) and had a range of gross pathologic diagnoses including multiple phenotypes of cerebrovascular disease. The pathologic Braak score is provided along with the post-mortem interval (PMI; hours). Sections of frontal peri-ventricular white matter were selected and underwent CXCL5 immunohistochemistry. Endothelial CXCL5 staining was graded in several ways. CXCL5 staining in any vessel within frontal peri-ventricular white matter was graded as positive CXCL5 staining (any positive vessel). The percentage of vessels with positive CXCL5 staining was also measured per total vessel segments visible in the section. Across this cohort, the majority of cerebral vessels in white matter displayed at least some CXCL5 staining within endothelia.

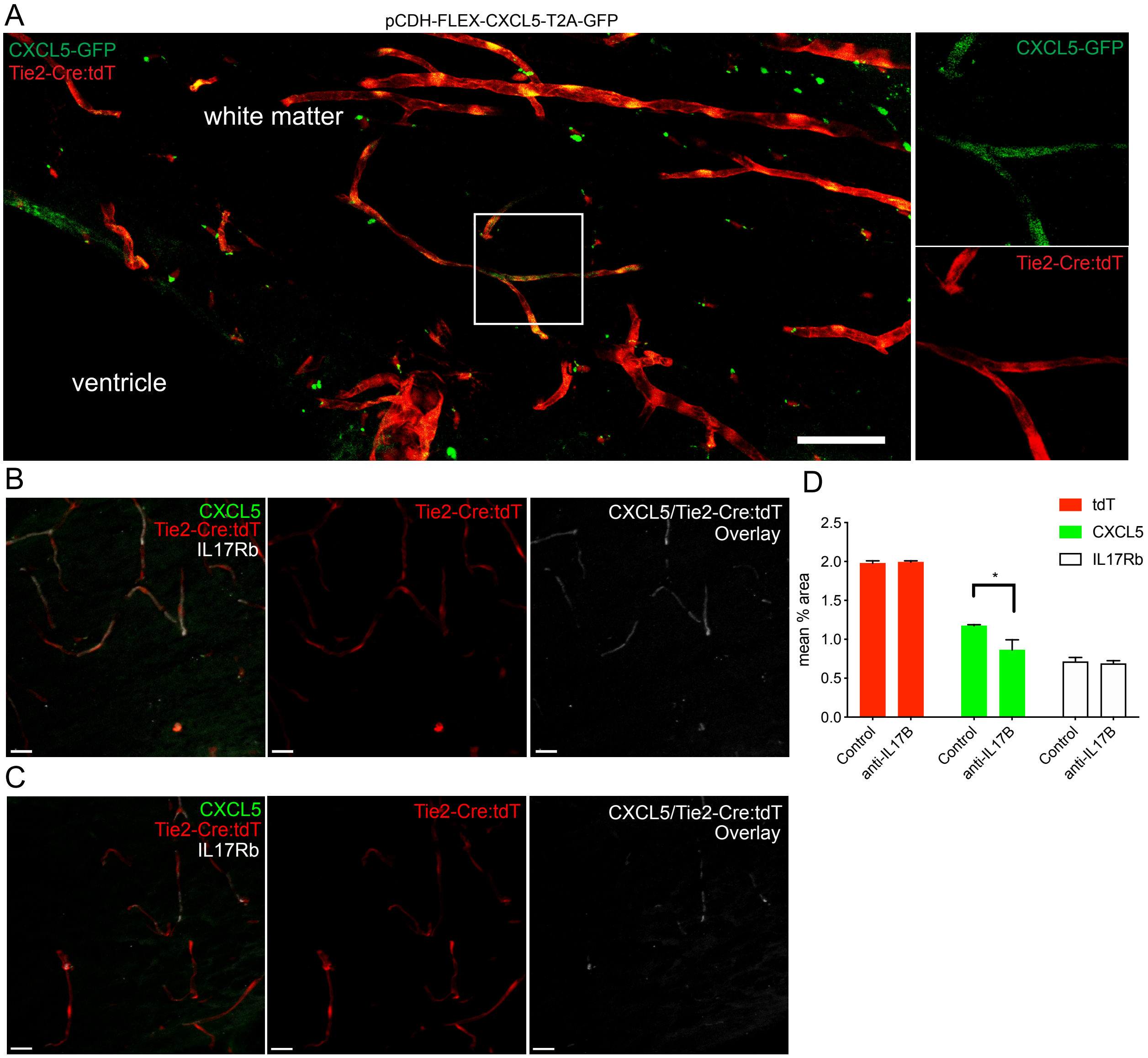


**Supplemental Figure 5. Modulation of CXCL5 expression in white matter vasculature using combined transgenic and viral transduction or anti-IL-17B IgG.**

Animals on control fat diet underwent viral transduction into the white matter with pCDH-FLEX-CXCL5-T2A-GFP lentivirus (shown) or control pCDH-FLEX-GFP lentivirus. Viral particles at a titer of $10^8$ were injected to white matter in Tie2-Cre;tdTomato mice at 14 weeks old and analyzed at 20 weeks of age. After 6 weeks of viral transduction, CXCL5-T2A-GFP (green) was highly colocalized with Tie2-Cre;tdTomato+ vessels (red) within the region of viral targeting in white matter (insets, A). To reduce DIO-induced endothelial expression of CXCL5, function blocking anti-IL-17B IgG or control isotype-matched IgG (50 µg) was administered q3d for 6 weeks in Tie2-Cre;tdTomato animals on HFD beginning at 14 months of age. At 20 weeks of age, immunolabeling for CXCL5 (green) and IL17Rb (white) was performed in animals receiving control IgG (B) or anti-IL-17B IgG (C). In white matter tdT+ vasculature, both CXCL5 and IL17Rb were colocalized with Tie2-Cre;tdTomato+ vessels. Compared to animals administered control IgG, mean % area occupied by immunolabeling of CXCL5 was significantly reduced in animals administered anti-IL-17B IgG (1.17 vs. 0.86; *$p$=0.0067; $n$=4) (D). Endothelial expression of IL17Rb (mean % area) was not significantly different between control IgG and anti-IL-17B IgG indicating that receptor expression was not altered by anti-IL17B IgG administration. Scale bars = 50 µm (A); 20 µm (B-C). Data are mean ± SEM.

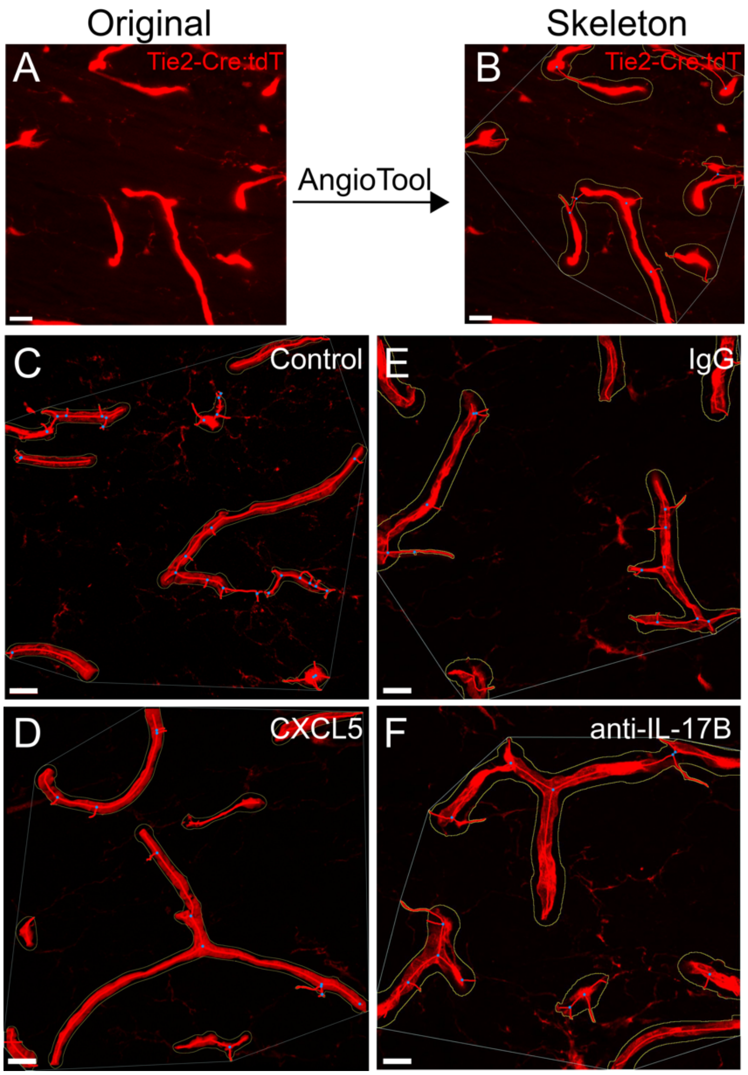


| Treatment | Vessel Length (mm) | SEM | *P-value* |
|---|---|---|---|
| Control (GFP) | 0.257216686 | 0.053521 | 0.891901056 |
| CXCL5-GFP | 0.253844777 | 0.05808 | |
| IgG | 0.218490871 | 0.037066 | 0.504302562 |
| anti-IL-17B | 0.207364 | 0.040578 | |

**Supplemental Figure 6. Measurement of microvascular length using AngioTool.**

The total vessel length of Tie2-Cre;tdTomato+ vessels were analyzed by AngioTool. The original high magnification tdT+ image from callosal white matter (A) was skeletonized in AngioTool which preserves contiguous vessel segments (outlines) and identifies branch points/junctions (blue dots) (B). Representative AngioTool vessel skeletons from animals on Tie2-Cre:FLEX-GFP expressing animals (C), Tie2-Cre:FLEX-CXCL5-T2A-GFP expressing animals (D), and animals on high fat diet administered either control IgG for 6 weeks (E) or anti-IL-17B IgG for 6 weeks (F). Average total vessel length of each treatment analyzed by AngioTool was shown in table. Data are mean ± SEM; Scale bar = 20 μm.

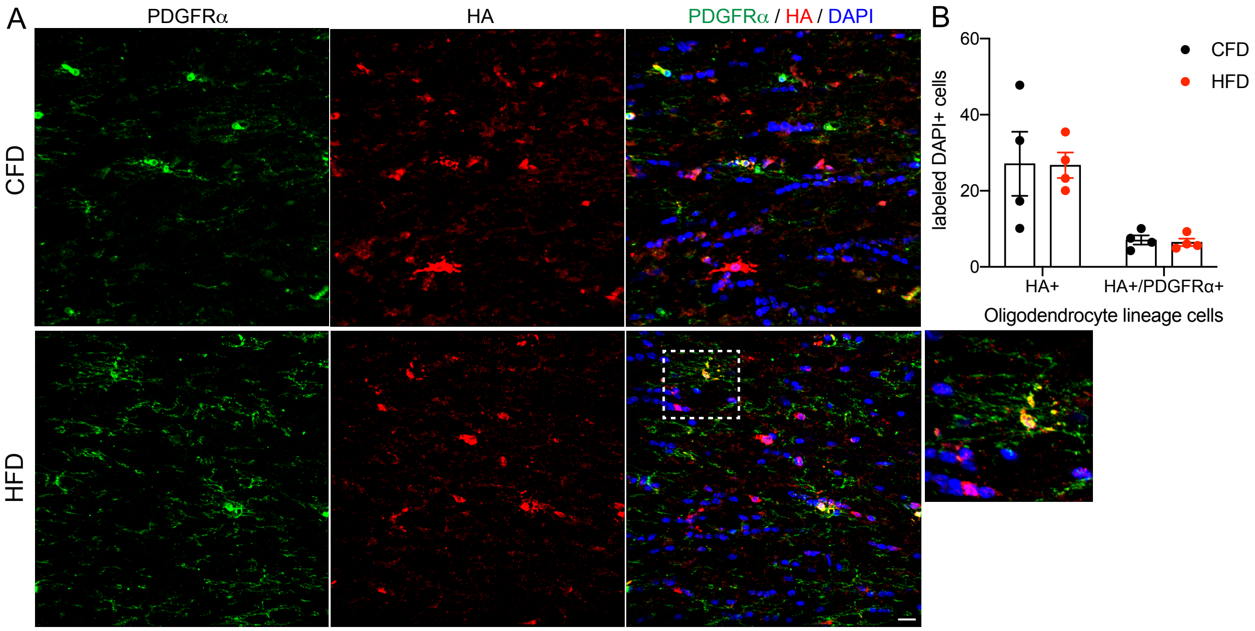


**Supplemental Figure 7. PDGFRα+ OPC fate mapping after onset of diet-induced obesity.**

After 20 weeks on a high-fat diet, PDGFRα-CreERT:RiboTag mice were injected with tamoxifen for 48 hrs and continued on control fat diet (CFD) or high fat diet (HFD) for 1 additional month (31.3±1.5g vs. 45.5±3.8g; $p$=0.014, $n$=4/grp). At 5 months of age, HA+ cells were fate mapped in callosal white matter from animals on CFD (upper panels) vs. HFD (lower panels) (A). HA+/DAPI+ cells (differentiated oligodendrocytes) and PDGFRα+/HA+/DAPI+ cells (OPCs) are distinct (inset) and the numbers of each respective cell lineage were counted in callosal white matter (B). PDGFRα+/HA+ (7.1±0.8 vs. 6.5±0.7) and HA+ (27.1±8.4 vs. 26.7±6.5) cells per 60X field of view were not significantly changed between animals on CFD vs. HFD, respectively ($p$=0.98 by 2-way ANOVA). Scale bar = 20 µm. Data are mean ± SEM.

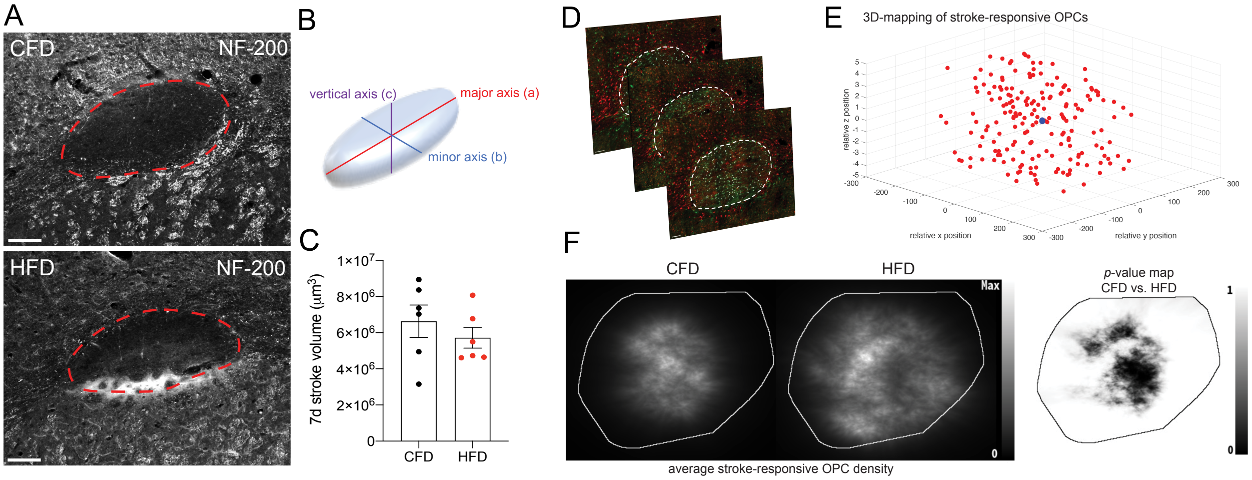


**Supplemental Figure 8. Effect of diet-induced obesity on white matter stroke volume.**

White matter stroke was induced as described producing a focal ischemic lesion precisely localized to subcortical white matter. The region of focal ischemia 7d post-stroke was identified by the absence of NF-200 axonal staining (A). Using serial sections, the elliptical volume of the stroke lesion was determined 7d post-stroke using formula $\pi/6 \times major\ axis \times vertical\ axis \times minor\ axis$ (B). Graph of white matter stroke volume 7d post-stroke between animals on CFD vs. HFD ($6.6 \times 10^6 \pm 604.0$ vs. $5.7 \times 10^6 \pm 485.8$ µm$^3$, $p$=0.31 by Mann-Whitney, $n$=6/grp) (C). Confocal z-stacks through the white matter stroke lesion were used to identify stroke-responsive PDGFRα+ OPCs (green) within the lesion core where GST-π+ mature oligodendrocytes (red) were lost due to focal ischemia (11 steps of Z-stacks per image) (D). Spatial coordinates of each cell (*x,y,z* in µm) relative to the center point of the stroke lesion (blue dot) were used to generate spatial maps of stroke-responsive OPCs (red) within each lesion section (9 sections/group; 3028 cells spatially measured). Representative coordinate map of stroke-responsive OPC localization from one section (E). Average OPC spatial density maps were generated in 2D for animals on CFD (left, $n$=3) and those on HFD (right, $n$=3) as described (J. Burguet et al Pattern Recogn Lett, 2011). Vertical gray bar represents cellular density with max density value = $3.97 \times 10^{-3}$ cells/µm$^2$. Curve in white: ROI contour. Map of $p$-values for differences in stroke-responsive OPC densities between animals on CFD and HFD. Vertical gray bar represents $p$-value. Low $p$-values (<0.05; black) correspond to regions with significantly more cells in the control group, and high $p$-values (>0.95; white) to regions with significantly more cells in the HFD group (F). Scale bar = 100 µm (A); 50 µm (D). Data are mean ± SEM.
